# Both environmental conditions and intra- and interspecific interactions influence the movements of a marine predator

**DOI:** 10.1101/2024.02.26.582077

**Authors:** Johann Mourier, Marc Soria, Matthew Silk, Angélique Demichelis, Laurent Dagorn, Tarek Hattab

**Affiliations:** MARBEC, Univ Montpellier, CNRS, IFREMER, IRD, Sète, France; Centre for Ecology and Conservation, University of Exeter Penryn Campus, Cornwall, UK; NIMBioS, The University of Tennessee Knoxville, Knoxville, TN, USA; CEFE, Univ Montpellier, CNRS, EPHE, IRD, Montpellier, France

**Keywords:** spatial-social interface, acoustic telemetry, *Carcharhinus leucas*, ERGM, movement networks, Indian Ocean, Reunion Island, shark

## Abstract

Animal movements are typically influenced by multiple environmental factors simultaneously and individuals vary in their response to this environmental heterogeneity. Therefore, understanding how environmental aspects, including biotic, abiotic and anthropogenic factors, influence the movements of wild animals is an important focus of wildlife research and conservation. We apply Exponential Random Graph Models (ERGMs) to analyse movement networks of a bull shark population in a network of acoustic receivers and identify the effects of environmental, social or other types of covariates on their movements. We found that intra- and interspecific factors often had stronger effects on movements than environmental variables. ERGMs proved to be a potentially useful tool for studying animal movement network data especially in the context of spatial attribute heterogeneity.

---

Understanding animal movement is not straightforward, as movement decisions result from a combination of many factors including the internal state of the individual, its intra- and interspecific neighbourhood, and spatio-temporal variation in abiotic environmental conditions (Nathan et al., 2008). Additionally, the response of individuals to variation in their surrounding environment can differ across spatial and temporal scales, as well as along their ontogeny. Daily activities can be influenced by environmental factors that vary over a diel cycle (e.g., wind or tidal currents), while seasonal movement can be governed by changes acting at larger temporal scales (e.g., photoperiod, temperature). Quantifying how heterogeneity in an animal’s surroundings drives behaviour, and consequently movement patterns, provides insight into how natural and anthropogenic changes may impact populations and ecosystems.

One potential way to understand the ‘causes, mechanisms and spatiotemporal patterns of movement and their role in various ecological and evolutionary processes’ (Nathan et al. 2008) is to link observed movement patterns with spatial and temporal variability in the underlying environmental conditions (Avgar et al., 2013). Movement patterns result from interactions between organisms and their environments (Johnson et al., 1992; Morales et al., 2010; Schick et al., 2008). For example, movement rates are expected to decrease where resources are rich and increase where food availability is low (Kuefler et al., 2012; Pyke et al., 1977). Understanding movement patterns thus requires consideration of the temporally dynamic nature of these environments (Avgar et al., 2013; Couriot et al., 2018; Mueller et al., 2011; Riotte-Lambert & Matthiopoulos, 2020; Schick et al., 2008).

In addition to interacting with their physical environment, animals also interact with other individuals of the same or different species. These spatial encounters with conspecific or heterospecific individuals may be advantageous or unfavourable. For example, aggregations form to avoid predation or to forage efficiently (Krause & Ruxton, 2002), although the tendency for non-gregarious species to seek or avoid each other is less well known. For all sexual species, individuals need to encounter conspecifics at least for mating, but avoiding opposite-sex individuals (sexual segregation) may play a major role in preventing sexual harassment in some species (Wearmouth & Sims, 2008). Additionally, avoidance strategies can act at the intra- or interspecific level in the context of competition for mates, space or resources (Giuggioli & Kenkre, 2014), or in the context of predator–prey interactions (Atwood et al., 2009). Because individuals may select habitats based on exogenous environmental factors (e.g. habitat quality and predation risk), and local population factors (e.g., mating opportunities, competition or density dependence), we need to quantify the mechanisms by which dynamic social interactions between individuals occur (e.g. movement towards or away from other individuals). In fact, interactions with conspecifics are intrinsically connected with spatial behaviour and a burgeoning literature now addresses the link between spatial and social aspects of behaviour defined as the “spatial-social interface” (Webber et al., 2023). While spatial and social processes have been predominantly considered independently, because movement behaviour emerges from social and spatial processes, animal movement studies offer an opportunity to consider and integrate them (Albery et al., 2021; He et al., 2019; Mourier et al., 2019; Webber et al., 2023).

Finally, the human footprint has altered the spatial ecology of many species at different spatial and temporal scales, for example by decreasing animal movements as a result of behavioural changes, habitat fragmentation and barrier effects (Doherty et al., 2021; Tucker et al., 2018), or by modifying activity-timing (Gilbert et al., 2023). COVID-19 lockdowns provided an empirical experiment where an abrupt reduction in human activity (so called Anthropause) led to decreases in displacements and a reduced avoidance of human activity (Tucker et al., 2023). Human disturbance can also fundamentally alter the way that species interact. For example, it can induce a spatiotemporal compression of species co-occurrences in disturbed landscape which can lead to increases in competition, predation and infectious disease transmission (Gilbert et al., 2022). As such, incorporating human footprint as a potential factor affecting animal movements at the spatial-social interface is now more relevant than ever.

In this study, we explore the connection between seascape attributes and animal movement patterns for a population of bull shark (*Carcharhinus leucas*) that inhabit a coastal zone in Reunion Island (Indian Ocean), which encompasses a variety of environmental conditions. We evaluate the relative importance of spatiotemporal variables associated with several main aspects of the seascape (abiotic conditions, social environment, predation risk or interspecific competition and anthropogenic pressure) as potential drivers of shark movement patterns. This approach allows us to identify fundamental relationships between local environmental conditions and animal movement patterns.

The bull shark is a large Carcharhinidae species with a wide cosmopolitan distribution along the continental coasts of all tropical and sub-tropical waters of the world, and is known to be mobile and move across a large range of habitats and environmental conditions (Brunnschweiler et al., 2010; Daly et al., 2014; Espinoza et al., 2016; Heupel et al., 2015; Lee et al., 2019; Niella et al., 2022). Additionally, it is also an euryhaline elasmobranch that uses a range of salinities throughout its life-cycle (Niella et al., 2022). Its behaviour varies across spatial and temporal scales, as well as according to size and sex, with a high among-individual variability in the tendency to move (Espinoza et al., 2016; Lee et al., 2019; Mourier, Soria, et al., 2021). However, much remains unclear about adult bull shark movement decisions. Bull shark movements are known to be influenced by both biotic and abiotic factors (Lee et al., 2019; Lubitz et al., 2023; Niella et al., 2022; Werry et al., 2018), and anthropogenic factors (Hammerschlag et al., 2022; Werry et al., 2012), but among the biotic factors, less is known about the importance of con- and heterospecifics in movement decisions. Even if most studies on the movements of these marine predators have identified rainfall, sea surface temperature and food distribution as the most influential to their movement ecology, we expect that social and competition factors may be just as important in explaining individual movements. Our analysis was designed to test whether individual bull sharks move according to the distribution of conspecifics and heterospecifics in the seascape, and whether these effects were comparable with expected positive effects of swell height, turbidity and rainfall on shark movements and negative effects of human densities (with sharks avoiding human presence). While we predict that bull sharks will respond to certain abiotic factors, being attracted to high levels of turbidity, rainfall and swell as revealed in other studies, we also expect the distribution of conspecifics and heterospecifics in the seascape may drive movement decisions and that male and female sharks may have different strategies.

## METHODS

### Study location

Reunion Island (21°07’S / 55°32’E) is a volcanic island located 700 km east of Madagascar in the southwest Indian Ocean. The island is 2512 km^2^ with 217 km of coastline and characterized by steep underwater slopes (*ca.* 10-20%) to a depth of 2,000 m. Fringing reefs stretch over 25 km along the west and south-west coast (Figure 1) forming a natural coral barrier that bounds the reef flats and back-reef depressions and lies no further than 500 m from the beach.

**Figure 1:**
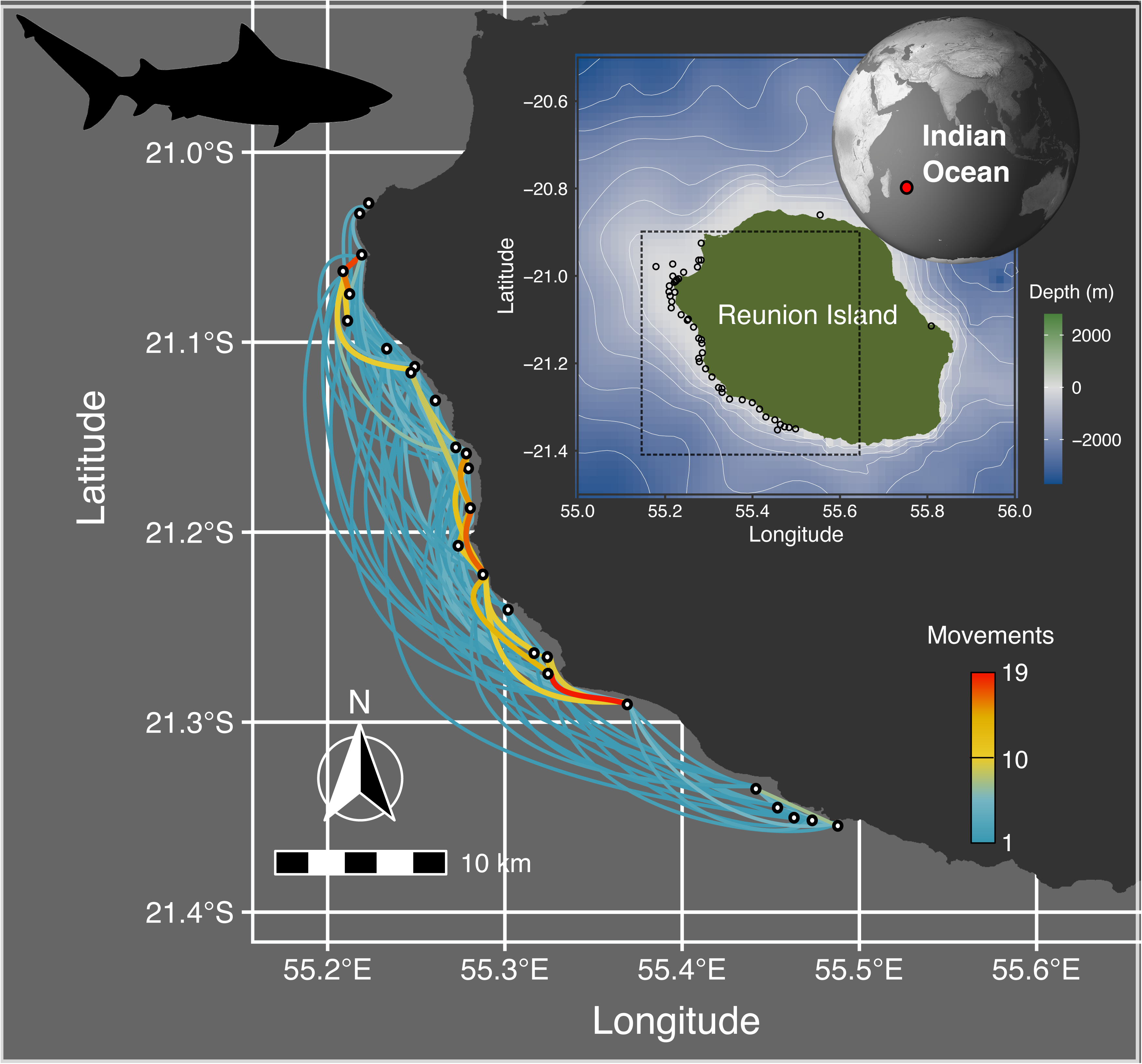
**C**umulative number of movements across all 44 monthly individual movements used in the ERGM analyses concentrated on the western part of Reunion Island. Circles represent acoustic receivers. Inset map represents the location of Reunion Island and the area where the movements were recorded.

### Acoustic telemetry and movement network

Sharks were captured along the west coast of Reunion Island between September 2012 and March 2013 (Supplementary material 1 Table S1), using horizontal drifting long-lines 0.2 to 1 km in length and equipped with 20 to 200 baited 16/0 circle hooks (Blaison et al., 2015). Set times were fixed at a maximum of 3 h to minimize shark and bycatch mortality. Once captured, a shark was brought alongside the vessel and held still by rubber-encased ropes to prevent skin lesions and burns and rolled onto its back to induce tonic immobility. The boat moved forward slowly throughout the procedure to allow the shark to breathe. Sanitized transmitters (Innovasea V16, transmission interval 40–80 s, estimated battery life 845 days) were implanted into the peritoneal cavity through a midventral incision using a sterile scalpel. A 1cm incision was made, then enlarged with retractors. Once the tag was in place, two stitches closed the incision and facilitated healing. A sterile, non-absorbable synthetic monofilament suture (polyamide) was used. The shark was then freed by cutting the hook at its base and removing it, untying the lasso, replacing it on its belly and oxygenating it by moving it back and forth or slowly forward until the first signs of autonomous movement appeared. Sex and total length (TL) were recorded during the procedure. A total of 23 individual sharks (i.e., 6 males and 17 females) were equipped with transmitters (Supplementary materials 1 Table S1).

An array of 46 Innovasea VR2W acoustic receivers was deployed along the coast with receivers installed an average of approximately 2 km apart at depths of 10–60 m, comprising 33 (71%) offshore receivers deployed between 300 m and 5 km from shore, and 13 (29%) inshore receivers placed less than 300 m from shore (Figure 1).

For each shark visit at a receiver, we used detection records to calculate a continuous residency time (CRT) corresponding to the duration within which a tagged shark was continuously monitored at a specific receiver (Capello et al., 2015; Ohta & Kakuma, 2005). All detections of the same shark at one receiver separated by less than a predefined period, called the maximum blanking period (1 h), were grouped into one CRT and defined as a visit. Each time a tagged shark was detected at a different receiver, a new visit started - ending the visit at the previous receiver - even if the interval between detections was less than the maximum blanking period. We then built monthly movement networks of each shark with each node representing a receiver along the coast of Reunion Island and each weighted, directed edge represented the number of movements of the individual (deduced from CRT) from node A to node B (Figure 1) within a given month.

### Explanatory variables

We gathered a variety of biological, abiotic, anthropogenic and spatial data that were accessible in the study area. Although non-exhaustive, the explanatory variables recorded included a number of environmental, biological and anthropogenic factors (Table 1; a detailed description of how they were recorded can be found in Supplementary material 1) as well as the geographical distances between receivers. We incorporated swell height, turbidity and rainfall levels as abiotic explanatory variables of movements. Turtle densities were included as a potential indicator of the presence of sharks, as suggested by local people.

**Table 1:** Predictor parameters included in the Exponential Random Graph Models for bull sharks monitored along the west coast of Reunion Island.

| Variables | Category | Description | Range | Source |
| --- | --- | --- | --- | --- |
| Occupancy Tiger sharks | Biological | Frequency of days at least a tiger shark was detected at the receiver for each month (%) | [0-0.409] | Current acoustic telemetry data |
| Same sex CRT | Biological | Total CRT of bull sharks of the same sex for each month (hour) | [0.033-118.425] | Current acoustic telemetry data |
| Opposite sex CRT | Biological | Total CRT of bull sharks of the opposite sex for each month (hour) | [0.033-118.425] | Current acoustic telemetry data |
| Bull Shark abundance | Biological | Number of bull sharks present at the listening station divided by the number of bull shark with an active tag for each month | [0-1.24] | Current acoustic telemetry data |
| Turtle density | Biological | Mean density of turtles in the area of the receiver from aerial survey (nb/km<sup>2</sup>) | [0-15.873] | Aerial surveys |
| Turbidity | Environmental | Average monthly reflectance for each zone over the multiple images (%) | [4.101-8.886] | Satellite SPOT 4 & 5 |
| Rain | Environmental | Mean rain fall for each watershed for each month (mm) | [0.008-25.460] | METEO-France |
| Swell | Environmental | Height of the swell near the listening station for each month (m) | [0.557-2.636] | AVISO portal, CANDHIS & METEOLAMER platform |
| Human activities | Anthropogenic | Cumulated number of human activities (boats, swimmers, nautical activities) for each month | [0-1276.5] | Aerial surveys |
| Geodist | Spatial | Distance between stations (km) | [0.9-120] | Current acoustic telemetry data |

Cumulative number of sea users (boats, swimmer and nautical activities) represents an anthropogenic variable. We also included various intra- and interspecific factors such as residency times of the opposite sex, abundance of bull sharks and occupancy of tiger sharks (a larger and likely competitor species) as con- and heterospecific social interaction variables. As all data were not available at the same spatial and temporal scale, we chose to standardize all available candidate explanatory data at the scale of one month for each receiver, which practically corresponded to calculating a mean for each explanatory variable at each receiver for each month. This monthly resolution represents a compromise between biologically meaningful variation in site conditions (i.e., node attributes) and retaining enough movements to construct individual networks. For some environmental variables such as rainfall, multiple nearby receivers shared values as data were available for a specific zone encompassing several receivers. All explanatory variables were standardized to have a mean of 0 and a standard deviation of 1.

### Exponential Random Graph Models

Care is required when conducting statistical analysis of network data because of issues related to potential non-independence among neighbouring nodes and edges (Croft et al., 2011). Thus, we used Exponential Random Graph Models (ERGMs). ERGMs are statistical models of networks that treat the weight of network edges as the response variable and network node and edge attributes as explanatory variables (Robins, Pattison, et al., 2007; Robins, Snijders, et al., 2007; Snijders et al., 2006). Such models are analogous to Generalized Linear Models (GLMs) except that they enable hypothesis testing about the processes driving network structure and link formation. These models have been mostly applied in social science, but their properties also make them useful for answering questions related to how and why animals move between locations in the context of movement networks (Fletcher et al., 2011; Jacoby & Freeman, 2016; López-Calderón et al., 2023).

Effectively the movement network becomes the response in a regression model, where the predictors are the propensity for nodes (i.e. locations) of similar or dissimilar attributes to be linked by movements. In the current study, we applied ERGMs to our monthly individual bull shark movement networks. These ERGMs are treat the weight of network edges (i.e. number of movements between two nodes) as the response variable and network node (i.e. acoustic receivers) attributes as explanatory variables.

The general form for an ERGM can be written as:

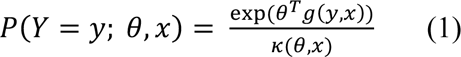

where:

- *Y* is the random variable for the state of the network (with realization *y*),
- *g(y,x)* is a vector of model statistics for network *y*,
- *θ* is the vector of coefficients for those statistics, and
- *κ(θ)* is a normalizing term which ensures that equation (1) is a proper probability distribution. It represents the quantity in the numerator summed over all possible networks (typically constrained to be all networks with the same node set as *y*).

The numerator represents a formula that is linear in the log form (equation 2):

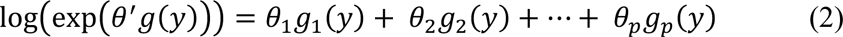

where *p* is the number of terms in the model. From this one we can more easily observe the analogy to a traditional statistical model. The functions *g(y)* are counts of configurations in the network *y* and the parameters *θ* weight the relative importance of the respective configurations, effectively the size and direction of the effects of the covariates. Parameter estimation in most specifications of ERGMs uses maximum pseudo-likelihood, an approximation of maximum likelihood based on Monte Carlo estimation.

All ERGMs were fitted using R packages *ergm* and *ergm.count* (Hunter et al., 2008).

### Model-fitting process

Node attributes were varying at the month scale so we built a model for each monthly-individual network. To ensure that it was possible to fit ERGMs to monthly networks and improve model convergence, we first removed from monthly movement networks all receivers that were deployed less than 20 days during the month, as well as all individuals that did not present an active tag for at least 20 days during a month. This empirical choice was made as a compromise between data loss and data representativity. We also retained only monthly individual networks containing at least 10 different directed movements between receivers, as networks including low numbers of movements caused problems with model convergence and parameter estimation. We also excluded two monthly networks that were binary (i.e. only contained at most single movements along any particular edge). This left us with 42 monthly networks of 13 individuals (9 females: 4 males; mean/median/range of monthly networks per individual = 3.1/3/1-6).

For each monthly network and each shark, we then fitted two alternative versions of the full model (using terms ‘nodecov’ and ‘absdiff’) accounting for different effects of the variables being studied (Table 1). The effect estimated for each factor represents the difference from the intercept as with classical GLMs. All models included the term “non-zero” to control for zero-inflation generated by the weighted movement networks being sparse. The term “sum”, corresponding to the sum of all link values, was included as the equivalent to the intercept in a linear model (Dey & Quinn, 2014).

The two model sets differed as follows:

- Model set 1: We tested the association between the current state of the environment at each receiver location and number of movements towards and away from it (i.e. effect of an attribute on out- and in-degree). These models used the *nodecov* term in the fitted ERGMs.
- Model set 2: We tested the association between the absolute difference in current state of the environment at dyads (pairs) of receiver locations and number of movements between them. These models used the *absdiff* term in the fitted ERGMs.

We used a *nodecov* model rather than separate *nodeicov* and *nodeocov* models (movements towards and away from receivers respectively) as simulations indicated the results were qualitatively identical (see Supplementary material 2).

We fitted the *nodecov* and *absdiff* models separately to facilitate model parsimony and convergence and test the robustness of the movement patterns discovered while controlling for the full suite of environmental variables. Indeed, incorporating all variables and model sets combinations would prevent convergence of model and render interpretation of output complex. To account for the effect of the spatial distribution of nodes in our models, we included as a covariate the matrix of distances between nodes.

For each model set, we then wrote a model formula (equation 3) including all potential predictors as follows:

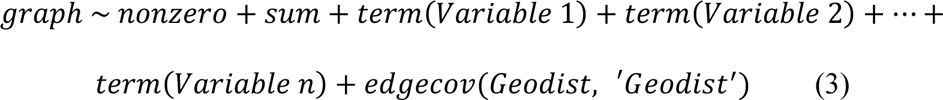

where *term* can be replaced by *nodecov* and *absdiff* in model sets 1 and 2, respectively. An *edgecov* term was added to control for the distance between locations. Models were fitted with a Poisson reference distribution for edge weights creating the familiar log-linear effect. The final coefficient estimates indicate log-linear increase in the weight variable (movements). MCMLE.Hummel.maxit was set to 1000 (Monte Carlo maximum likelihood estimation using the Partial Stepping technique of <u>Hummel et al., (2012)</u>) and MPLE.type was set to “penalized” so that the Maximum Penalized Likelihood Estimate(MPLE) was calculated using a bias-reduced method.

For each model set for each individual monthly network, we used Aikaike Information Criterion (AIC) to compare the AIC of 49 candidate models nested within the full model, and keep only the model that most parsimoniously explained the movements of the individual during the month considered. Any models with convergence issues were discarded.

We then re-fitted all selected ERGMs for each combination of both model sets (*nodecov* and *absdiff*) and individual monthly networks. From these final models, we extracted model estimates and their standard errors, and also MCMC diagnostics (Gewecke’s convergence diagnostic and *R̂*) to ensure convergence.

To summarize the results incorporating each selected models, we extracted parameter estimates and associated standard errors from selected variables from each selected models, and followed a meta-analysis procedure in which each sample (monthly-individual movement network) was treated as a single ‘study’. Effect sizes were calculated using an inverse-variance weighting meta-analysis for each sample and grouped by variables using a sub-grouping analysis. This allows to test if differences in effect sizes exist only due to sampling error, or because of true differences in the effect sizes. This procedure was made for the overall samples, as well as separating by sex and by season. The meta-analysis was conducted using the package ‘*meta*’ in R (Schwarzer, 2007).

### Simulation methods

To ensure the correct interpretation of our model results and check for potential limitations of the ERGM methods applied to our dataset, we used a simulation approach. In short, we simulated a series of individual movement trajectories that matched our empirically collected data. We then fitted ERGMs in the same way we did for the empirical data.

We generated sets of 25 receiver locations (equivalent to the empirical data) positioned in 2D space and generated three environmental properties for each receiver location termed factorA, factorB and factorC. We then generated 21 simulation input parameter sets that varied the effect of factorA on the probability of movements between locations while keeping factorB and factorC fixed as having no effects on movement (Supplementary material 2 Table S2). We considered scenarios where movements were conducted according to a gradient in factorA or occurred between similarly high or low factorA locations. For each simulation input parameter set, we simulated the movements of 10 sharks. The number of movements for each shark was drawn from a Poisson distribution with a mean of 23 (equivalent to the empirical dataset). Detailed methods are in Supplementary material 2.

## RESULTS

### Interpretations of model outputs from simulations

Our simulation study revealed ERGMs performed as expected when analysing movement networks of a similar size and structure to our empirically-measured networks (see Supplementary materials 2). Model estimates for the *nodecov* and *absdiff* models typically accurately represented the presence of movements up a gradient or between similar locations, respectively (Figure 2). The statistical power to detect true effects was limited for individual networks, and there were slightly elevated false positive rates for the *absdiff* models.

**Figure 2:**
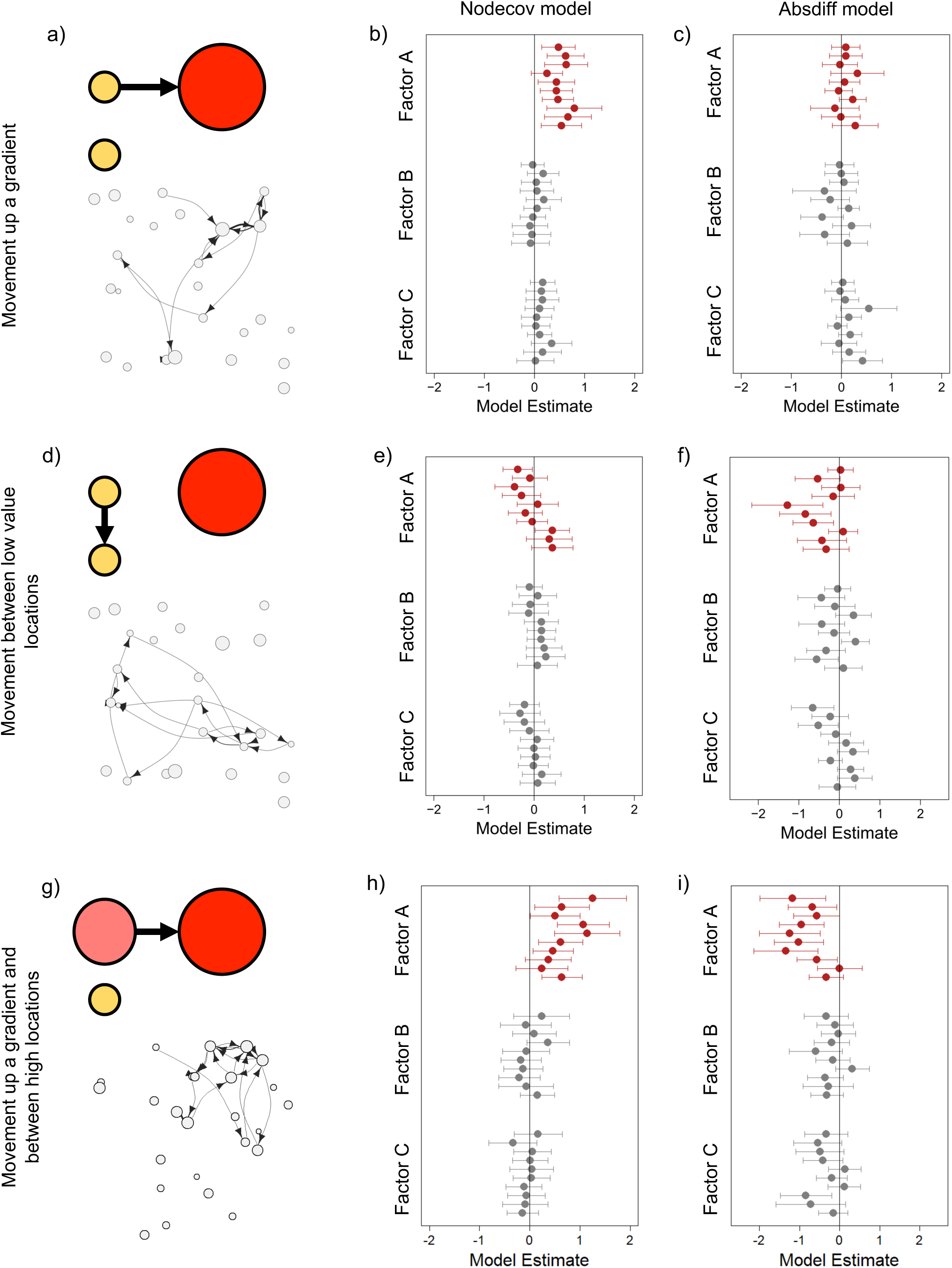
Interpretation of the model output for each scenario based on simulations: (a, b, c) movements up a gradient of FactorA (e.g. swell height), (d, e, f) movements between locations of low FactorA and (g, h, i) movement up a gradient and between locations of high FactorA. (a, d, g) Toy examples on the left and movement network for the first shark. (b, e, h) model estimates for *nodecov* and (c, f, i) model estimates for *absdiff*. The model estimates are for the three effects (highlighting the one where there is a true effect in red) and then for each of the 10 sharks simulated.

However, effect size estimates were largely unbiased (both with and without true simulated effects). The only exception was that *absdiff* models underestimated the tendency to move between similar locations when there was also a gradient effect (Figure 2f). In addition, *absdiff* models overestimated the tendency to move between similar locations when sharks tended to start moving from high value locations and had a strong tendency to move up a gradient (Figure 2i, although the latter would be expected if this resulted in sharks moving only between high value locations). Collectively, these results indicate that collating estimates from multiple models (as we do in our main analysis) will provide the most informative results. Simulations also indicated that *nodeicov* and *nodeocov* models provided closely correlated results leading to us using a single *nodecov* model in our empirical analysis (see above).

### Effects of abiotic conditions

Our models were designed to test whether sharks were moving between locations with similar abiotic conditions (tested by the *absdiff* term with a negative coefficient indicating movement between location with similar conditions, while a positive coefficient means that an increase in the absolute differences of abiotic conditions increases the odds of movements) and/or towards locations with higher/lower abiotic conditions (tested by the *nodecov* term, a positive coefficient indicating higher values and a negative coefficient suggesting lower values), specifically swell height, degree of turbidity and level of rainfall. Model results indicated the environmental variables that were associated with movement network structure (Figure 3). While high swell at a location was not significantly associated with more movements (positive coefficients in the *nodecov* model), movements between locations that differed considerably in swell height were much less frequent than movements between locations with similar swells as demonstrated by negative values of the *absdiff* model.

**Figure 3:**
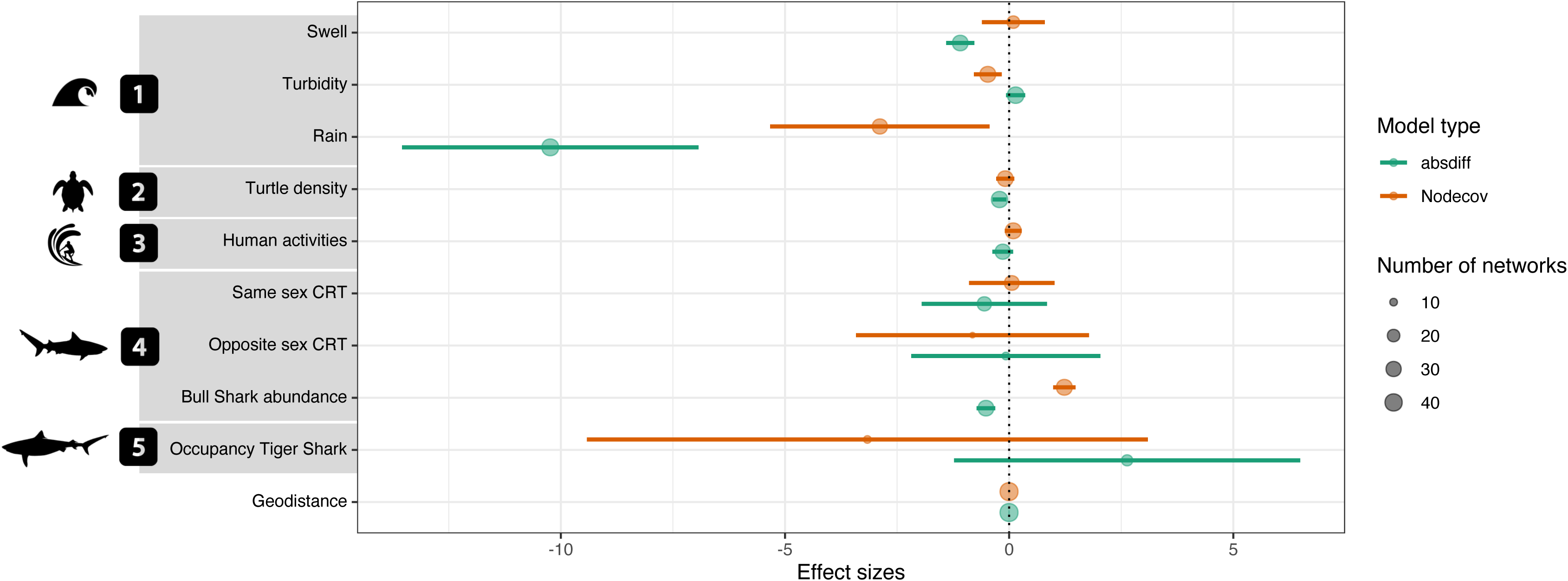
Summary of the outputs from the Exponential Random Graph Models from the 42 selected individual monthly movement networks. Effect sizes and their 95% confidence intervals pooled over variables are reported for the parameter estimate of terms *nodecov* and *absdiff*. Variables were grouped in several categories: (1) environmental factors, (2) turtle density, (3) anthropogenic factors, (4) social or intraspecific interactions, and (5) competitive or interspecific interactions. The variable “Geodistance” controls for the distance between receivers. Circle size is proportional to the number of models in which the coefficient of the term was significant and therefore selected.

Together, these outputs indicate that sharks move between locations characterized by similar swell and not across a gradient of swell heights. The *nodecov* model revealed a negative effect of turbidity on movements, indicating that more movements occurred through low turbidity locations. Further, the *absdiff* model revealed that sharks were also more likely to move between receiver locations that differed from each other in their turbidity. Collectively these results suggest that low turbidity increased the frequency of movements between receivers. Sharks also tended to move between receiver locations with similar and low rainfall levels. The observed trends were more robustly represented for rain and turbidity than swell as these variables were supported by more monthly networks and involved in a larger number of individuals with statistically significant results (Supplementary material 1 Table S2).

### Effect of turtle densities

Our models could also test whether sharks were moving toward high or low densities of turtles (tested by the *nodecov* term) and/or were remaining in habitats with similar densities (tested by the *absdiff* term). Model outputs indicated that movements were more likely between receiver locations with more similar turtle densities (negative estimate from *absdiff* model). However, parameter estimates were small and the number of monthly networks and individuals in which the coefficients were statistically significant were low (Supplementary material 1 Table S2), indicating that the biological importance of turtle distribution densities may be limited.

### Effect of anthropogenic factors

Our models were also set up to test whether sharks were avoiding (tested by the *nodecov* term with an expected negative coefficient) and/or remaining in areas of similar human activity (tested by the *absdiff* term with an expected negative coefficient). Models indicated that there were more movements among locations with high human activities (positive *nodecov* effect and negative *absdiff* effect). However, parameter estimates from the models were close to zero and the number of monthly networks and individuals in which the coefficients were statistically significant were relatively high (Supplementary material 1 Table S2), suggesting that the biological importance of anthropogenic factors such as human activities on bull shark movements in the monitored area were relatively limited but consistent among sharks and months.

### Influence of conspecifics

Our models were also used to test whether sharks moved towards or avoided conspecifics (tested by the *nodecov* term) and/or remained within areas with similar populations of conspecifics (tested by the *absdiff* term). Receiver locations in which bull sharks were present or more abundant were more strongly connected in the movement network. Outputs from *absdiff* model also indicated that movements tended to occur between locations with similar bull shark abundance, suggesting individuals were predominantly moving among a subset of preferred locations. While controlling for the abundance effect, there was (overall) some evidence for a negative effect of the residency time of sharks of the opposite sex, suggesting some spatial segregation between the sexes, but this effect varied seasonally (see below). The results were as robust, as the models describing responses to abiotic factors, characterized by a relatively high number of monthly networks and individuals involved and consistency in the direction of responses (positive or negative coefficients) across networks (Supplementary material 1 Table S2).

### Effect of inter-specific competition

We also used our models to test whether the presence of larger tiger sharks was influencing bull shark movements by moving to/away from areas used by tiger sharks (tested by the *nodecov* term) and/or moving between sites with similar numbers of tiger sharks (tested by the *absdiff* term). There was weak evidence (due to wide confidence intervals and the small number of times this parameter was selected in the top model, Supplementary material 1 Table S2) for the presence of tiger sharks affecting bull shark movement network structure. Positive effects in *absdiff* model indicated individuals tended to move across gradients of tiger shark occupancy, perhaps indicating active avoidance of tiger sharks, with some additional weak evidence of reduced movements through locations with higher occupancy of tiger sharks.

### Sex differences in movement network structure

Analysing movement network structure for the two sexes independently revealed similar broad trends as the overall movement network, except in the case of social factors and the tiger shark occupancy (Figure 4). While females tended to move toward locations with high residency times for other females, males showed a general pattern of avoiding locations with high residency times of either sex (while controlling for overall bull shark abundance). The weak overall evidence for reduced movements through locations with high tiger shark occupancy was driven by divergent effects between females and males. Female movements were directed towards locations with low tiger shark occupancy while male movements were directed towards locations with high tiger shark occupancy, indicating sex differences in how competitors influenced movement through receiver locations, although these effects have high uncertainty around them.

**Figure 4:**
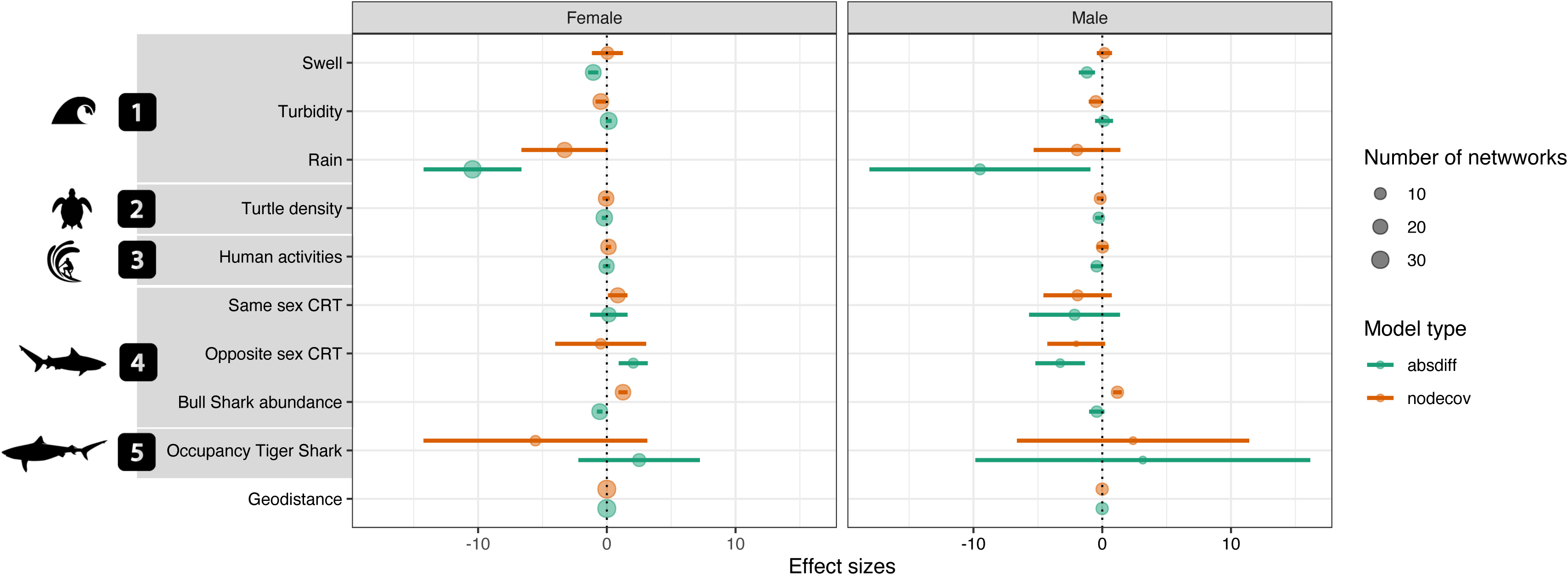
Summary of the outputs from the Exponential Random Graph Models presented separately for females (n = 10) and males (n = 3). Effect sizes and their 95% confidence intervals pooled over variables are reported for the coefficient of terms *nodecov* and *absdiff*. The variable “Geodistance” controls for the distance between receivers. Circles represent the number of networks in which the parameter estimate of the term was statistically significant and therefore selected.

### Seasonal differences in movement network structure

The factors that explained movement network structure remained similar between winter and summer, with only interspecific competition and rainfall changing qualitatively (Figure 5).

**Figure 5:**
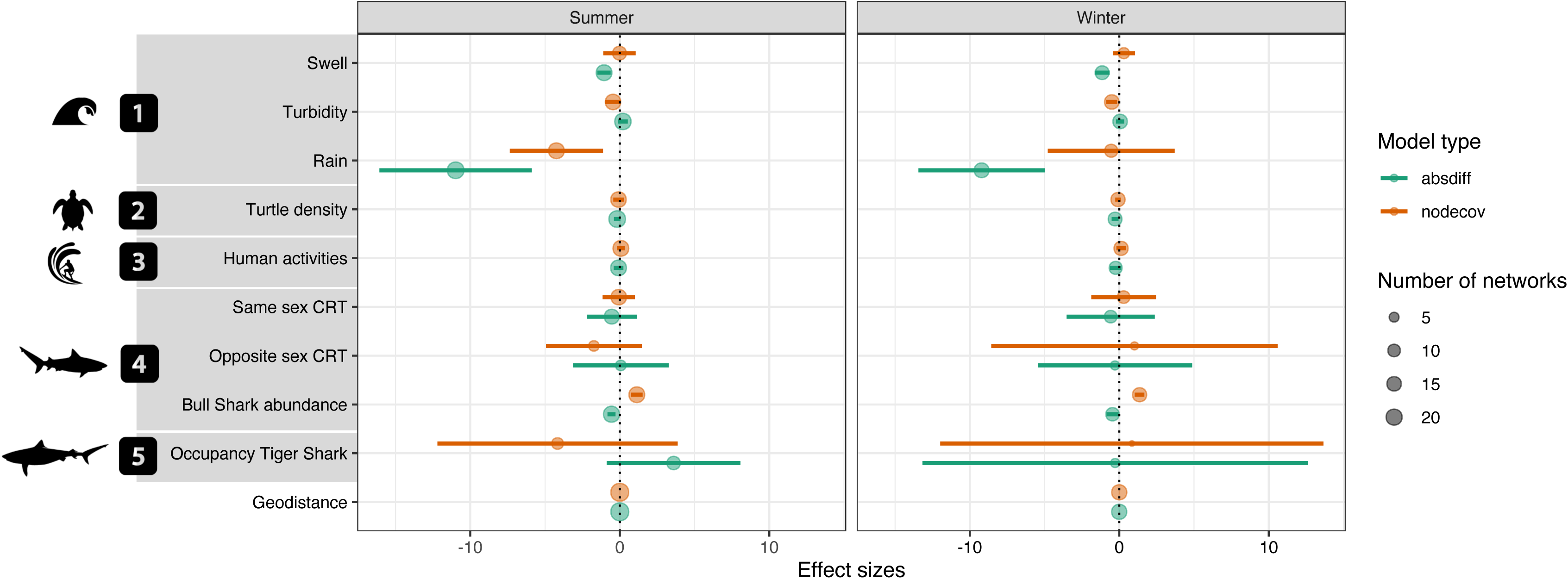
Summary of the outputs from the Exponential Random Graph Models presented separately for summer and winter. Effect sizes and their 95% confidence intervals pooled over variables are reported for the coefficient of terms *nodecov* and *absdiff*. The variable “Geodistance” controls for the distance between receivers. Circles represent the number of networks in which the coefficient of the term was significant and therefore selected.

While rainfall levels had effects on movements in both seasons, the pattern was less clear during winter, which is expected as rainfall is a seasonally-driven factor. It appears there is a shift in behaviour related to the presence of tiger sharks. While there was no apparent movement pattern related to tiger shark occupancy during winter, there was a pattern of movements towards locations with low occupancy of tiger shark in summer as demonstrated by positive *absdiff* coefficients and negatives coefficient of *nodecov*.

## DISCUSSION

We predicted that bull sharks would be attracted to high levels of turbidity, rainfall and swell. We also expected that bull shark movements would be influenced by the distribution of conspecifics and heterospecifics in the seascape, although we did not predict the direction of these effects. We were thus interested whether these con- and heterospecific effects were as important as abiotic variables in explaining movement patterns. As we hypothesized, if bull shark movements were driven by abiotic parameters, our models revealed that social factors were also important factors even if the reproducibility and consistency across the sampled individuals and months were slightly smaller (Supplementary material 1 Table S2). We also acknowledge that other potential important factors such as temperature or prey distribution not included in our model could contribute to explain shark movement. Additionally, the direction of response to abiotic variables were not always as predicted, indicating that the role of physical variables in shaping movement network structure may be context dependent or vary across spatial and temporal scales.

### Effects of abiotic factors on movements

Our results indicate that environmental factors impact the movements of bull sharks, however, while we expected to find positive effects of swell height, turbidity and rainfall on shark movements as suggested by previous studies, we found opposite patterns. Bull sharks were found to be mostly moving through areas with lower rainfall. While bull sharks are known to respond to rainfall and freshwater runoff and are able to transit between freshwater and marine habitats within hours or remain in low salinity areas for days (Niella et al., 2022; Werry et al., 2018), selection of lower rainfall levels could be due to the spatial and temporal resolution at which they react to such rapid environmental changes. In addition, bull sharks may prefer to predominantly remain (at our scale of observation) in the same conditions of rain, turbidity and salinity as rain levels are often strong and fast in tropical areas. Werry et al. (2018) suggested that bull sharks are attracted to estuarine and associated nearshore areas following high rainfall events in part due to increased prey availability while Niella et al. (2022) suggested a more complex pattern with differences in response between the sexes.

Matich et al., (2020) also highlighted that during exceptional events bull sharks can avoid high rainfall and high turbidity, moving away from sites with fast and strong perturbations. The rainfall data entered in our models represent distinct watershed values that can potentially incorporate multiple receivers, and can potentially explain why our models found disproportionately strong effects of movements between (nearby) locations with similar levels of rainfall. In addition, Werry et al. (2018) found a one-week lag in the response of sharks to rainfall which could reflect changes in food availability and foraging effectiveness with changes in salinity, a pattern that would be difficult to reveal with our models. Previous studies that found strong behavioural responses of bull sharks to rainfall were conducted in large estuarine system where rainfall levels and consequences (e.g. induced turbidity) could contrast with the coral reef coast of Reunion Island. Finally, movements in response to attractive effects of environmental factors (swell, turbidity and rainfall) that are ephemeral and irregular may occur over shorter timescales and be difficult to detect with monthly data. While we expected to find stronger seasonal differences because the variations in these parameters are more important and persistent in summer than during winter, the lack of such behavioural differences could be due to bull sharks being resident for less lengthy periods during summer (Blaison et al., 2015). It is important to note that other potential factors that we did not measure could also contribute significantly to movement decisions. For example, salinity is known to be an important factor affecting movement and distribution of bull sharks (Dwyer et al., 2020). Our results confirm that considering temporally dynamic environmental variables in studies of animal movement is important because movement patterns reflect dynamic interactions between animals and their physical environment, with environmental heterogeneity driving movements of individuals and allowing them to remain in optimal environmental conditions (Avgar et al., 2013; Mueller et al., 2011; Schick et al., 2008).

### Effects of turtle density on movements

Our study did not find any clear effect of turtle density on bull shark movements. While we used available data on turtle density to investigate its influence on movement patterns of bull sharks in this study based on local people’s perception, we acknowledge that turtles are not the main prey of bull sharks and remain opportunistic items (Cliff & Dudley, 1991). Bull sharks are known to feed mainly on teleost fish (Trystram et al., 2016) and future studies could integrate dynamic species-specific fishery data or parallel underwater surveys to investigate the role of abundance and composition of fish communities in driving predator movements. Indeed, resource availability and heterogeneity are important drivers of animal movement, especially for predators (i.e. prey abundance and distribution). Finding prey requires a predator to make adaptive decisions about which movement patterns to adopt to feed most profitably given a particular prey distribution (Riotte-Lambert & Matthiopoulos, 2020; Sims et al., 2006). Thus, interpreting predator movements within a prey landscape may provide a clearer picture of why certain habitats are selected over others.

### Effects of human densities on movements

While we detected a tendency for more movements through locations with high human use, the effect sizes were small suggesting human activities did not strongly influence movement patterns. This is not surprising as nautical human activities and density are spread fairly evenly along the west coast of the island where the network of receivers was deployed. In addition, it is now recognized that bull sharks can adapt to urbanized areas and do not especially avoid these high human density areas and their activities (Hammerschlag et al., 2022; Werry et al., 2012). The majority of large-bodied terrestrial carnivores tend to avoid high human densities and activities (Tucker et al., 2018, 2023). COVID lockdowns revealed the same trend in the marine realm with fish communities responding positively to the absence of human activities (e.g., fish biomass increasing and predatory species increasing usage of shallow habitat during tourism lockdown; Weng et al., 2023). However, it is less clear with sharks as they have previously been shown to both respond positively to cessation of human activities (Afonso, 2024) and adapt to high human densities (Hammerschlag et al., 2022). The limited suitable habitat available around the island may also prevent bull sharks from avoiding most human activities along the coast. In addition, human activities are concentrated on the west coast of the island and overlap with other factors that could explain bull shark habitat preferences, such as the presence of a marine protected area (Soria et al., 2019). Further investigations are therefore necessary to better understand the influence of human activities on bull shark behaviour.

### Effects of social environment on movements

We found that social factors sometimes had similar effects on movement networks to abiotic variables. While the bull sharks have been found to form occasional aggregations around fish farms (Loiseau et al., 2016) or at artificial provisioning sites (Bouveroux et al., 2021), they are not typically recognized as displaying collective behaviours. In addition, strong patterns of sexual and ontogenetic spatial segregation have been found in our study population (Mourier et al., 2021). However, we found that more movements occurred through bull shark hotspots. When focusing on the behaviour of each sex, we showed that males were less social and tended to avoid other sharks regardless of their sex. Contrastingly, females’ movements occurred between locations where other females spent a lot of time but avoided areas heavily used by males. This confirmed previous findings that females show stronger patterns of residency providing the opportunity to co-occur with other females, while males favour roaming behaviour (Mourier et al., 2021). This gregarious behaviour of females could also suggest the presence of mating arenas where females gather to choose transient males allowing them to avoid male harassment during the mating period. Further research investigating the spatio-temporal interactions between bull sharks is required to better understand their aggregative and avoidance behaviour. Our results thus confirm that an individual’s movements are not solely driven by abiotic factors. Individuals share space with other conspecifics, linking spatial and social processes (Albery et al., 2021; Webber et al., 2023). For example, the spatial distribution of resources inherently drives the proximity of individuals through foraging aggregations, influencing many forms of social interaction (Macdonald, 1983). As such, the distribution of individuals of a species in space generates population structure and will influence movement decisions at the individual level. In fact, regardless of whether a species is perceived to be solitary or social, individuals will have to decide whether to join or avoid other conspecifics distributed unevenly in space. Such decisions will also depend on social processes driven by individual phenotypes that alter social decisions (e.g. size, sex, or genetic relatedness).

### Effects of interspecific competition on movements

In our study we tested the influence of another large predator, the tiger shark, on the movements of bull sharks. Our results indicated the presence of patterns of avoidance between bull and tiger sharks. Bull shark movements were affected by the presence of tiger sharks, with female bull sharks directing their movements towards locations with lower tiger shark occupancy while male sharks showed the opposite pattern. The tiger shark is an apex predator that is larger than the bull shark and potentially dominant. It is therefore plausible that bull sharks avoid interactions with tiger sharks. Female bull sharks, being larger than males (Hoarau et al., 2021), could also compete with tiger sharks. Therefore, one plausible explanation to this pattern is spatial and foraging niche separation between both species (Niella et al., 2021). We previously highlighted that male and female bull sharks displayed patterns of spatial segregation (Mourier et al., 2021; Niella et al., 2022), which could also explain the difference in interaction opportunities with tiger sharks based on habitat utilization and movement patterns. These patterns of interference interactions are relatively common in large sharks, as competitive species generally show spatial (Papastamatiou et al., 2018), temporal (Lear et al., 2021; Séguigne et al., 2023) and trophic niche (Matich et al., 2017) partitioning to limit the negative effect of competition and promote co-existence of predators. Similar avoidance strategies are found between competiting carnivores in terrestrial ecosystems. Indeed, lions (*Panthera leo*) usually remain in areas rich in prey, while subordinated carnivores like leopard (*P. pardus*) and cheetah (*Acinonyx jubatus*) overlap with the home range of lions but use fine-scaled avoidance behaviours, and small species like African wild dog (*Lycaon pictus*) employ multiple tactics to avoid all other competitors (Vanak et al., 2013). Coexistence of multiple carnivore species is typically explained by dietary niche separation as a consequence of avoiding intraguild competition. Another potential explanation is the avoidance of tiger sharks by females during summer corresponding to the parturition season (Pirog et al., 2019) in order to avoid tiger shark predation on bull shark neonates.

Although our results are to be treated with caution because we only studied a small number of individuals, and only a subset of these individuals responded to tiger sharks (Table 2), the direction of response was consistent. Our results thus complement previous studies suggesting that interactions among large predators involve a complex interplay of competition and predation, as large carnivores can suppress populations of smaller carnivores through direct predation, resource competition, or via other forms of interference competition. This may result in spatial and/or temporal avoidance, reductions in the density of the subordinate species, or even competitive exclusion from certain habitats (Berger & Gese, 2007; Linnell & Strand, 2000; Prugh et al., 2009; Prugh & Sivy, 2020).

### Conclusion

To investigate the drivers of movements of a marine predator, we used a network-based approach of movement between fixed stations and employed Exponential Random Graph Model in a way that could account for the dynamical nature of site attributes visited by individual animals within a flexible framework that could be extended to test hypotheses related to the structure of the network itself (extending beyond what is easily possible using more conventional statistical approaches). While this modelling framework performed well in our study, it could be refined further to test similar research questions at different spatial and temporal scales. For example, one promising model is the separable temporal exponential-family random graph model (tERGM), which treats the formation and dissolution of ties in parallel at each time step as independent ERGMs (Carnegie et al., 2015; Krivitsky & Handcock, 2014) and can provide a more realistic view of movement networks (e.g., including consecutive individual monthly or daily networks). This would make it possible to better integrate the temporal interdependency of environmental changes in explaining animal movement patterns.

Our results fit into a broader picture illustrating that animal movements arise from complex interactions of individuals with their physical environment as well as with both surrounding conspecific and heterospecific individuals with various level of consistency. By including a diverse set of variables that may influence bull shark movements within our analyses, we demonstrate that to fully explain animal movement patterns requires the incorporation of multiple variables associated with environmental heterogeneity, human footprint and the distribution of individuals in space, both of the same species and other members of the community. A main finding was that, although environmental conditions were important factors influencing movement of bull sharks, interactions with other individuals in their surroundings was also important to consider, opening new perspectives at the socio-spatial interface for these marine predators.

## Author contributions

**Johann Mourier**: Conceptualization (Equal); Formal analysis (Equal); Investigation (Equal); Methodology (Equal); Software (Equal); Validation (Equal); Visualization (Equal); Writing – original draft (Lead); Writing – review & editing (Equal). **Marc Soria**: Conceptualization (Equal); Data curation (Equal); Funding acquisition (Equal); Investigation (Equal); Methodology (Equal); Project administration (Equal); Resources (Equal); Supervision (Equal); Validation (Equal); Writing – review & editing (Equal). **Matthew Silk**: Formal analysis (Equal); Investigation (Equal); Methodology (Equal); Software (Equal); Validation (Equal); Visualization (Equal); Writing – review & editing (Equal). **Angelique Demichelis**: Formal analysis (Equal); Investigation (Equal); Software (Equal); Writing – review & editing (Equal). **Laurent Dagorn**: Conceptualization (Equal); Investigation (Equal); Validation (Equal); Writing – review & editing (Equal). **Tarek Hattab**: Conceptualization (Equal); Investigation (Equal); Methodology (Equal); Supervision (Equal); Validation (Equal); Writing – review & editing (Equal).

## Acknowledgments

We are grateful to the members of the institutions and associations involved in the CHARC program (Connaissance de l’HAbitat des Requins Côtiers de la Réunion): IRD, CRPMEM, CRESSM, University of Reunion Island, Globice, Kélonia, ARVAM, Squal’Idées, RNMR, and IFREMER as well as the fishermen and volunteers who assisted with the shark tagging and made our work possible. This study received financial support from the European Union (convention FEDER ref. 2012-dossier Presage no. 33021), the Ministère de la Transition écologique et Solidaire (BOP 113 n°2012/03), and the Regional Council of Reunion Island (POLENV no. 20120257). MJS received funding from the European Union’s Horizon 2020 research and innovation programme under the Marie Sklodowska-Curie grant agreement No 101023948. All the fieldwork and protocols of handling and tagging of sharks were approved by the Ethics Committee (n° 114) for the CYROI (Cyclotron Réunion Océan Indien).

Procedures were adapted to minimize stress on animals and avoid mortality (more details in Supplementary material 1). All operations were carried out or supervised by scientists with a certificate in animal experimentation and a certificate in experimental surgery (Oniris, Ecole Nationale Vétérinaire de Nantes).

## Data availability

Data and R codes used in this study are available at SEANOE: https://doi.org/10.17882/99080

