## Supplementary Materials 1 for "Both environmental conditions and intra- and interspecific interactions influence the movements of a marine predator"

**Supplementary material 1**

Table S1: Summary of the 23 tagged bull sharks used in this study conducted from September 2012 to May 2014. For each individual shark are given the following information: name, sex, size in nearest centimetre, date of tagging, number of detections recorded by the receivers, location of tagging and release and number of networks used in this study the individual was represented.

|  | Shark code | Sex | Total length (cm) | Tagging date | No. of detections | Location of tagging and release | Lat. Release Position | Long. Release Position | No. of networks |
| --- | --- | --- | --- | --- | --- | --- | --- | --- | --- |
|  | Shark01 | F | 329 | 28/9/12 | 12409 | Saint-Gilles | -20.96398 | 55.15310 | 10 |
|  | Shark02 | F | 314 | 5/11/12 | 3869 | Saint-Gilles | -21.05658 | 55.20853 | 4 |
|  | Shark03 | F | 314 | 10/2/12 | 22179 | Saint-Gilles | -21.04485 | 55.18747 | 2 |
|  | Shark04 | F | 310 | 6/1/13 | 3784 | Saint-Paul | -20.99753 | 55.27510 | 8 |
|  | Shark05 | F | 310 | 27/9/12 | 127 | Saint-Gilles | -21.06643 | 55.21608 | 1 |
|  | Shark06 | F | 308 | 10/2/12 | 19952 | Saint-Gilles | -21.05307 | 55.22098 | 4 |
|  | Shark07 | F | 307 | 24/3/13 | 5117 | Saint-Gilles | -21.05319 | 55.20128 | 9 |
|  | Shark08 | F | 300 | 10/1/13 | 49 | Saint-Gilles | -21.05347 | 55.21312 | 1 |
|  | Shark09 | F | 300 | 10/2/12 | 6593 | Saint-Gilles | -21.05293 | 55.22113 | 2 |
|  | Shark10 | F | 300 | 25/2/14 | 2546 | Saint-Gilles | -21.01130 | 55.17630 | 3 |
|  | Shark11 | F | 300 | 27/2/13 | 6889 | Saint-Gilles | -21.06333 | 55.21483 | 7 |
|  | Shark12 | F | 300 | 6/2/13 | 2123 | Saint-Gilles | -21.04878 | 55.20680 | 8 |
|  | Shark13 | F | 300 | 10/1/13 | 164 | Saint-Gilles | -21.05267 | 55.21350 | 1 |
|  | Shark14 | M | 300 | 6/1/13 | 2089 | Saint-Paul | -20.98950 | 55.27750 | 2 |
|  | Shark15 | M | 294 | 26/3/13 | 7644 | Saint-Pierre | -21.34662 | 55.43228 | 11 |
|  | Shark16 | M | 290 | 15/3/13 | 1128 | Saint-Pierre | -21.34380 | 55.43675 | 5 |
|  | Shark17 | F | 274 | 28/9/12 | 12573 | Saint-Gilles | -21.03200 | 55.21982 | 20 |
|  | Shark18 | M | 269 | 1/3/13 | 5251 | Etang-Salé | -21.23700 | 55.29300 | 11 |
|  | Shark19 | M | 260 | 19/3/13 | 7599 | Saint-Gilles | -21.04332 | 55.19787 | 12 |
|  | Shark20 | M | 260 | 20/2/13 | 15356 | Saint-Pierre | -21.34258 | 55.43528 | 14 |
|  | Shark21 | F | 260 | 20/2/13 | 7003 | Saint-Pierre | -21.34000 | 55.43175 | 14 |
|  | Shark22 | F | 238 | 6/1/13 | 15451 | Saint-Paul | -20.93313 | 55.27800 | 17 |
|  | Shark23 | F | 183 | 29/11/12 | 43218 | Saint-Louis | -21.30840 | 55.38320 | 15 |

Table S2: Summary of the statistically significant models providing for each variable the number of monthly networks with significant coefficients, the number of negative and positive coefficients, and the number of individuals in which each coefficient was statistically significant including the number of males and females.

| Model | Variables | Monthly networks | Negative coefficients | Positive coefficients | Individuals (Male-Female) |
| --- | --- | --- | --- | --- | --- |
| nodecov | Swell | 6 | 1 | 5 | 5 (2M-3F) |
|  | Turbidity | 15 | 13 | 2 | 11 (3M-8F) |
|  | Rain | 12 | 9 | 3 | 8 (3M-5F) |
|  | Turtle density | 6 | 4 | 2 | 5 (1M-4F) |
|  | Human activities | 9 | 2 | 7 | 8 (3M-5F) |
|  | Same sex CRT | 6 | 3 | 3 | 6 (2M-4F) |
|  | Opposite sex CRT | 5 | 4 | 1 | 3 (1M-2F) |
|  | Bull shark abundance | 18 | 0 | 18 | 9 (3M-6F) |
|  | Tiger shark occupancy | 6 | 3 | 3 | 4 (1M-3F) |
| absdiff | Swell | 19 | 19 | 0 | 9 (2M-7F) |
|  | Turbidity | 8 | 1 | 7 | 5 (2M-3F) |
|  | Rain | 21 | 21 | 0 | 12 (2M-10F) |
|  | Turtle density | 4 | 2 | 2 | 3 (2M-1F) |
|  | Human activities | 7 | 4 | 3 | 6 (2M-4F) |
|  | Same sex CRT | 10 | 2 | 8 | 7 (3M-4F) |
|  | Opposite sex CRT | 4 | 1 | 3 | 4 (1M-3F) |
|  | Bull shark abundance | 9 | 8 | 1 | 7 (2M-5F) |
|  | Tiger shark occupancy | 5 | 0 | 5 | 4 (1M-3F) |

*Ethical Note*

All the fieldwork and protocols of handling and tagging of sharks were approved by the Ethics Committee (n° 114) for the CYROI (Cyclotron Réunion Océan Indien). Procedures were adapted to minimize stress on animals and avoid mortality. The use of circular hooks prevented the shark from swallowing them. All sharks swam away in good condition after being released. The procedure usually lasted less than 30 min. Most sharks were detected and therefore remained alive during the experiment (mean days of detections 88.3 ± 79.4; min = 1, max = 285). Two sharks were caught and removed by local fishermen and two others were never detected after being released. All operations were carried out or supervised by scientists with a certificate in animal experimentation and a certificate in experimental surgery (Oniris, Ecole Nationale Vétérinaire de Nantes).

*Environmental data acquisition*

Turbidity

The data come from SPOT 4 and 5 satellite images recorded by the SEAS-OI (Indian Ocean Satellite Aided Environment Monitoring antenna (http://www.seas-oi.org/)) receiving satellite image. Previous work showed that when the value of turbidity increases, the value of the reflectance of a certain wavelength increases. The wavelengths most sensitive to these variations are in the blue, green, then in red and near infrared (Ouillon et al., 2008). The wavelengths used are between 500 to 590 nm (Doxaran et al., 2002).

The preprocessing of the satellite images makes it possible to superimpose the images correctly and to correct them for the effects of the atmosphere in order to have a set of images that are comparable between them. The processing steps are as follows (ENVI 5.1 software):

• The first preprocessing consists in ortho-rectifying the image acquired at level 1A (http://www.astrium-geo.com/fr/911-niveaux-de-pretraitement-et-precision-de-localisation). The Digital Terrain Model used for this purpose is that of the IGN at 25 meters resolution. The images are then calibrated between them based on a SPOT 5 ortho-image (level 3) dated May 30, 2011 of the BD-Isle of Kalideos;

• Then in order to compare the images (Kergomard, 2000), a correction for the effects of the atmosphere (resulting from the gases and aerosols contained in the atmosphere leading to the creation of absorption and diffusion (Kergomard, 2000; Girard and Girard, 2010) effects) is applied to each of the images. This correction is applied to each referenced image (date, time, satellite, longitude / latitude, image gains) from data collection on AOT (Aerosol Optical Thickness) at 550 nm. The data comes from the global Aeronet network (http://aeronet.gsfc.nasa.gov/). The output file contains for each image: the spectral clear water reflectance, the solar spectrum, and the total gas transmittance (total gaseous transmittance). These three values make it possible to apply the correction calculations to the image studied and thus obtain an image corrected for the effects of the atmosphere and therefore comparable to other images having followed the same treatment.

Once these values have been acquired, the application of atmospheric corrections involves three calculations, the result of the first of which allows the calculation of the second, and thus until the last calculation. The details of each calculation are specified below, and appear in order of application:

◦ Calculation of luminance or radiance: Once the images are ortho-rectified and calibrated, they are calibrated in radiance value by applying the gain values indicated in the specifications of each image.

- $L=\frac{\mathrm{Cn}}{\mathrm{Ce}}$

Cn = Digital raw image count

Ce = Calibration coefficient

◦ Calculation of the exo-atmospheric reflectance:

- $Rea=100*\frac{\pi*L}{\cos\left( \theta\right)*Eˉ^{1}s}$

Cos(ϴ) = solar zenith angle

E^-1^ s= equivalent solar irradiance$=\frac{Solar spectrum}{Band width}$

◦ Calculation of ground reflectance:

- $Rs=\frac{Rea-R atm}{⅂g}$

R atm = Atmospheric reflectance requiring the value of the spectral reflectance in clear water

⅂g = Total gas transmission for the channel considered

The exo-atmospheric reflectance and ground reflectance calculations were applied only to the green channel of each image. To keep only useful information, the preprocessed images were vectorized in QGIS. Then a geographic mask was applied to the images making it possible to mask the emerged areas and reefs, finally a radiometric mask is applied to the marine parts in order to keep only the useful measurement interval. The uninformed areas (clouds) are characterized by a “no data” coding. The reflectance values are between 1 and 12 at sea and characterize the concept of water color associated with the concept of turbidity.

The study area was divided into 58 distinct zones from north to south depending on whether they are located along the coast (27 sectors of average area = 1.24 ± 0.38 km²), coastal (27 coastal sections of 3.23 ± 0.99 km²) or offshore around 4 DCP (Area = 3.8 km²). For each zone (Zoning from the CHARC program), on each image acquisition date, an average reflectance value, a maximum value, a minimum value and a standard deviation were calculated. This resulted in an attribute table, comprising 52 zones, for each of the images processed. From February 7, 2012 to November 21, 2013, 68 satellite images were analyzed.

Rain

METEO-France provided daily precipitation data (in mm) between 2011 and 2013 on around thirty stations of interest for the study area.

From this information, the stations were grouped according to their belonging to the different watersheds which, together with the altitude data, allow the zone to be categorized into different climatic regions (Robert, 1986). Ten climatic regions have been defined on the west coast of Reunion Island with between 1 to 7 stations per region. For each of these regions, the mean precipitation deviations between monthly and annual data were calculated and served as a baseline.

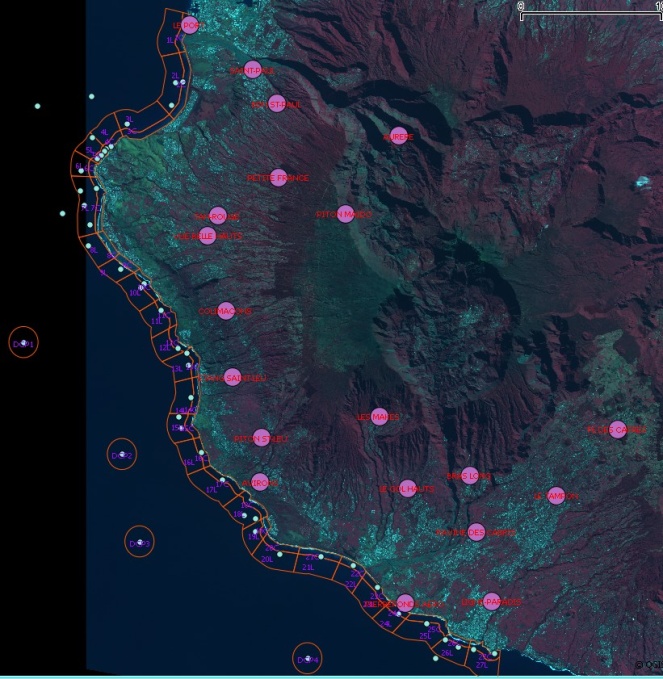

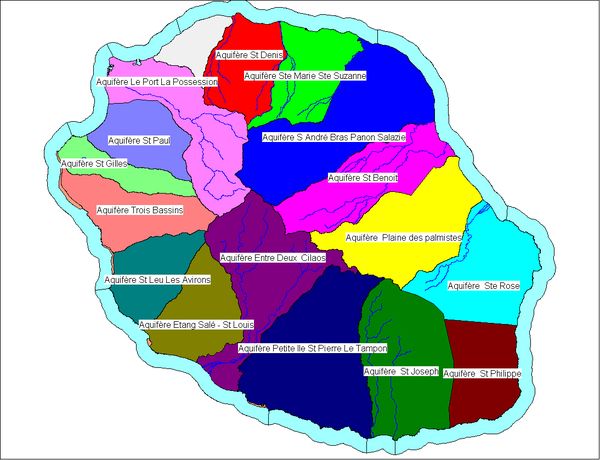

Figure S1: Climatic regions (left) resulting from stations grouped according to their belonging to the different watersheds (an example on the right).

Swell

Swell data were obtained from different sources: from the AVISO portal (Archiving, Validation and Interpretation of Oceanographic Satellite Data) for water level data, from CANDHIS (National Archiving Center for In Situ Swell Data) and from the METEOLAMER modeling platform. This platform provides a model making it possible to characterize the physical environment of Reunion Island at different scales.

The values were estimated every 6 hours at 12 sites around Reunion Island (StDenis, Houlographe, LePortOuest, StPaul, Boucan, RochesNoires, Ermitage, TroisBassins, StLeu, EtgSalé, StPierre and SteRose), between January 01, 2012 and January 01 2015.

On these sites are estimated: the significant height and the maximum height, the significant period and the maximum period of the waves defined by passage at the mean level by decreasing values (recommended by AIRH 1986).

From this data and the water depth at these sites, an orbital bottom speed value was estimated for each sample. The orbital speed of the particles Uw associated with the passage of the wave near the bottom (𝑚 s-1) was expressed according to the following formula:

U_w_ = (π x H/T) x (1/sin h [ 4 x π² x H/ (g x T²)])

where H is the height of the swell (m), T is the period of the swell (s), g is the gravity (ms-2), h is the height of the water column (m).

Aerial survey used to infer human activities and turtle density

West coast turtle surveys were taken from 34 ULM flights between Saint-Pierre and Le Port, from March 2012 to August 2013 at the rate of one flight every 15 days (2 flights per month). Each flight consisted of a one-hour round trip along the coastline to report and map ocean human activities including all kind of fishing, tourist or diving boats, nautical activity, surf, bathers and beach attendants. All kind of activities were cumulated.

Sea turtle count was carried out on the same area, from September 2012 to September 2013 during 41 flights (3-4 flights per month). The coastal strip flown over, along a sinusoidal transect, was about 1.5 km wide, and included the entire Marine Nature Reserve. The flights were conducted at an average altitude of 200m at an average speed of 90km per hour. The flights were carried out according to standard meteorological conditions (swell < 2 meters, winds < 9 knots and a clear sky) between 8:00 a.m. and 10:00 a.m. in order to reduce the impact of nautical activities on the presence of sea turtles and the time spent on the surface. Along this transect, the turtles present at the surface were counted and located using a GPS (Global Positioning System). Observation densities were calculated from the estimated area overflown per zone (54 CHARC meshes). This area flown over was estimated by multiplying the distance traveled within an area by the width of the "observation swath" estimated at 150 m on either side of the ULM (or 300 m in total).

Bull and tiger shark abundances

Abundance of bull and tiger sharks were calculated by summing the total number of individuals that visited each receiver for each month.

Bull and tiger shark occupancy

Occupancy was calculated as the percentage of days at least an individual bull or tiger shark, respectively, was detected at each receiver for each month.

Same and opposite sex bull shark Continuous Residency Time

This corresponded to the cumulated Continuous Residency Time in minutes of all sharks from the same or opposite sex of the shark considered, respectively.
