## Supplementary Materials 2 for "Both environmental conditions and intra- and interspecific interactions influence the movements of a marine predator"

**Supplementary material 2**

Simulation Methods

To ensure the correct interpretation of our model results and check for potential limitations of the ERGM methods applied to our dataset we used a simulation approach. In short, we simulated a series of movement trajectories for individuals that matched our empirically collected data but varied the effect size of parameters that influenced the probabilities of movement. We then fitted ERGMs in the same way we did for the empirical data and quantified their false positive (likelihood of an ERGM detecting a statistically significant effect when no effect was simulated) and false negative (likelihood of an ERGM not detecting a statistically significant effect when a true effect was simulated) error rates.

Movement network simulation

We simulated a simplified version of our empirical dataset. We generated sets of 25 receiver locations (equivalent to the empirical data) positioned in 2D space. The x and y coordinates of receiver locations were drawn from a uniform distribution. We then randomly generated three environmental properties for each receiver location factorA, factorB and factorC. Each environmental property was randomly simulated from a Normal distribution with a mean of zero and standard deviation of one.

We then generated 21 parameter sets (used as inputs to generate movement networks) that varied the effect of factorA on the probability of movements between locations while keeping the two other environmental variables fixed as having no effects on movement (Table S2). Either the likelihood or number of movements was increased to a location or locations with higher factorA (GRADIENT) or more movements were likely between sites with more similar values of factorA (DIFFERENCE). For each case we considered 7 parameter values increasing in strength from no effect to a strong effect. We considered both mechanisms acting alone and also the case where both factorA and similarity in factorA had an effect on the number of movements between locations.

*Table S2. Simulated effect size combinations.*

| *Parameter set* | *Gradient effect size* | *Difference effect size* |
| --- | --- | --- |
| *1* | *0* | *0* |
| *2* | *0* | *0.25* |
| *3* | *0* | *0.5* |
| *4* | *0* | *0.75* |
| *5* | *0* | *1* |
| *6* | *0* | *1.25* |
| *7* | *0* | *1.5* |
| *8* | *0* | *0* |
| *9* | *0.25* | *0* |
| *10* | *0.5* | *0* |
| *11* | *0.75* | *0* |
| *12* | *1* | *0* |
| *13* | *1.25* | *0* |
| *14* | *1.5* | *0* |
| *15* | *0* | *0* |
| *16* | *0.25* | *0.25* |
| *17* | *0.5* | *0.5* |
| *18* | *0.75* | *0.75* |
| *19* | *1* | *1* |
| *20* | *1.25* | *1.25* |
| *21* | *1.5* | *1.5* |

For each parameter set we simulated the movements of 10 sharks. We simulated scenarios with three different starting location tendencies. Either a) each shark started their movements at a randomly selected receiver location; b) starting location selection was biased so that sharks were more likely to begin at low factorA locations; or c) starting location selection was biased so that sharks were more likely to begin at high factorA locations. The number of movements for each shark was drawn from a Poisson distribution with a mean of 23 (equivalent to the empirical dataset). The probability of moving to a new location then depended on: a) how close other receiver locations were to the current location (weighted as the inverse of distance squared); b) how much higher factorAs were at potential new locations (movements more likely to locations with higher factorAs); and c) how similar a location the factorA was at potential new locations (movements more likely to locations with more similar factorAs). The importance of b) and c) were determined by the effect sizes in each parameter set (see previous paragraph). Simulation code is provided at <https://github.com/johannmourier/ERGMshark>.

In total we simulated 630 movement networks (10 sharks for each of 21 parameter sets and 3 starting location biases).

ERGM fitting

After simulating each set of 10 movement networks (1 per individual for each parameter set) we fitted three alternative ERGMs. This is similar to the main text but with the addition of one model. We fitted models containing either a nodeocov, nodeicov (movements away from and towards the receivers respectively) or absdiff term for each of our environmental variables (factorA, factorB and factorC), an edgecov term to control for the distance between locations. Models with either fitted a) using the R package ergm.count and having a Poisson reference distribution; or b) using the R package ergm to fit models to any unweighted networks. We used the latter only when there were not any repeat movements between the same pairs of locations. While there were some networks in the empirical dataset that were binary they were excluded from the final outputs due to our use of a meta-analysis type approach (model coefficients are not directly comparable between Poisson and Binomial implementations of the model). For ergm.count models we included the terms sum (a equivalent to an intercept in a GLM) and non-zero (to control for zero-inflation in the movements recorded in the network). For ergm models we included the term edges to control for zero-inflation. During the design of the simulations we encountered problems with model convergence in some runs of the simulation. Consequently, we included a timeout function for all ERGMs run during the simulations, with any model that took longer than 60 seconds to fit excluded. We then extracted the model coefficients from all successfully fitted models.

In total we attempted to fit 1890 ERGMs, three models for each simulated movement network.

Analysis

In the main text we present single scenarios (generated separately from the full simulation results) from: a) random start locations, move up gradient only; b) low factorA start locations, move between similar locations; and c) high factorA start locations, move up gradient and between similar locations. In each scenario the effect sizes are at there strongest (i.e. parameter combinations 7, 14 and 21 respectively). We use these illustrations to aid with model interpretation for the real results and demonstrate how the simulation approach worked.

To generate the full results, for each environmental variable (factorA, factorB and factorC) we considered the proportion of simulated sharks for which we detected a statistically significant effect for a given combination of variable and model in each simulation (combining results from both binary and weighted networks). If the models were working appropriately then we would expect to detect the patterns outlined in Table S3. Any deviations from these patterns indicate potential limitations of using ERGMs to model these datasets. We also examined the distribution of ERGM-estimated effect sizes (from weighted network models only) to help explain any unexpected patterns in model performance. Finally we compared the estimates of the effect of factorA from models containing the nodeicov() and nodeocov() terms to determine the need for fitting both models to the empirical data.

*Table S3. Expected performance of ERGMs applied to shark movement data under ideal performance.*

| Model term | Nodeocov model | Nodeicov model | Absdiff model |
| --- | --- | --- | --- |
| FactorA | For parameter sets 8-14 and 15-21 the number of statistically significant results should show a saturating positive relationship with the strength of the effect size  For parameter sets 1-7, consistently low levels of statistically significant results independent of the simulated effect sizes | For parameter sets 8-14 and 15-21 the number of statistically significant results should show a saturating positive relationship with the strength of the effect size  For parameter sets 1-7, consistently low levels of statistically significant results independent of the simulated effect sizes | For parameter sets 1-7 and 15-21 the number of statistically significant results should show a saturating positive relationship with the strength of the effect size  For parameter sets 8-14, consistently low levels of statistically significant results independent of the simulated effect sizes |
| FactorB | Consistently low levels of statistically significant results independent of the simulated effect sizes | Consistently low levels of statistically significant results independent of the simulated effect sizes | Consistently low levels of statistically significant results independent of the simulated effect sizes |
| FactorC | Consistently low levels of statistically significant results independent of the simulated effect sizes | Consistently low levels of statistically significant results independent of the simulated effect sizes | Consistently low levels of statistically significant results independent of the simulated effect sizes |

Simulation Results

Our simulation results (Figs. S1-S30) indicate that ERGMs are performing largely as expected when applied to studying individual movement networks. However, there are important limitations to performance to highlight that we therefore account for when using and interpreting model results.

***Random starting locations***

Detection rates

For both the nodeicov() models (Figs. S1-2) and nodeocov() models (Figs. S3-4) model performance was largely as predicted. As the true effect of factorA increased the likelihood of the models detecting a statistically significant relationship, although statistical power was limited even with a strong effect of factorA. The pattern was broadly equivalent when there was both only a gradient effect and when this gradient effect was combined with a difference effect. This is perhaps unsurprising given the networks are both small (low number of nodes) and sparse (low number of edges). When there was only an effect of the difference in factorA early there was a slightly elevated false positive rate but this remains at approximately 10% or lower regardless of the strength of this effect. False positive rates for the two variables with no effects on movement (factorB and factorC) remained close to 5% regardless of how factorA influenced movement patterns.

ERGM performance (in terms of detection rates) for the absdiff models (Figs. S5-6) was less impressive. The models were able to detect movements tending to occur between more similar locations when these occurred in isolation, although power was again limited. However, when a tendency to move between similar locations was combined with a tendency to move up a gradient power was very limited indeed. Further false positive rates were substantially elevated for factorA when no difference effect was simulated. There was a slight indication that the false positive rate for factorA increased when there was a strong gradient effect.

Effect sizes

For all models effect size distributions were broadly as expected (Figs. S8-10). When a gradient effect was simulated the nodeicov() and nodeocov() models estimated positive effects of factorA that increased with the value of the true effect size (Figs. S8-9). Effect size estimates were similar regardless of whether the gradient effect occurred in isolation or in combination with the difference effect. When a difference effect was simulated the absdiff() model estimated negative effects (as more movements occurred between sites with less different factorAs) that decreased with the true effect size (Fig. S10). However, the effect sizes estimated with the difference effect occurred alongside a gradient effect were smaller than when it occurred in isolation helping to explain the very low power in detecting these effects. In almost all cases when there was no true effect (both for factorA with confounders present and for other variables with no effect) parameter estimates were unbiased and centred on zero. The one exception was for absdiff() models when there was no difference effect and a strong gradient effect, in which case there was a very small upward bias in effect size estimates. Overall, these results indicate that both limited power and slightly elevated false positive rates are associated with small/sparse network effects rather than any biases in parameter estimates.

Relationship between nodeicov() and nodeocov() models

Estimates in nodeocov and nodeicov models were strongly correlated with the slope of the relationship very close to 1 indicating that using both of these models would be redundant (Fig. S7).

***Biased starting locations***

When we repeated the simulations with starting receivers biased towards low factorA locations (Figs. S11-20) or high factorA locations (Figs. S21-30) our results were qualitatively the same and quantitatively extremely similar to when starting locations were random, but we draw attention to some potentially important differences here.

Detection rates

When starting locations were biased towards low factorA receivers patterns were broadly similar but: a) there tended to be a slight increase in false positives for nodeicov() and nodeocov() models with strong difference effects simulated (Figs. S11, S13); b) false positive rates for the two variables with no effect were very slightly higher in all models (Figs. S12, S14, S16); c) there was a greater power to detect a tendency to move between similar locations when it was combined with a gradient effect (Fig. S15); d) false positive rates for factorA in the absdiff() model increased more substantially when there was a strong gradient effect.

When starting locations were biased towards high factorA receivers results for the nodeicov() and nodeocov() models were very similar to those for random starting locations (Figs. S21-24). For the abdsiff() model there was elevated power to detect a true difference effect when it occurred in isolation, but unlike the low-biased scenario a minimal change in power to detect a true difference effect in combination with a gradient effect (Fig. S25). Similarly to the low-biased scenario the increase in false positive rates with a strong gradient effect was more notable (Fig. S25) than when starting locations were random.

Effect sizes

Patterns for effect size distributions were extremely similar for both low-biased (Figs. S18-20) and high-biased (Figs. S28-30) starting locations. The only noteworthy patterns was that the small upward bias in effect size estimates for absdiff() models when there was no difference effect and a strong gradient effect for random starting locations was exacerbated in the high-biased scenario (Fig. S30) and not apparent in the low-biased scenario (Fig. S20).

Relationship between nodeicov() and nodeocov() models

The same strongly correlation between nodeicov() and nodeocov() estimates existed when starting locations were biased (Figs. S17 & S27). However, when starting locations were biased towards high factorA nodeicov() estimates tended to be slightly larger and when they were biased towards towards low factorA nodeocov() estimates tended to be slightly larger.

Simulation Discussion

There are a two key take-home messages from our simulation results:

1. Estimates derived from nodeicov() and nodeocov() models in small individual movements networks such as those in our studies are effectively equivalent and it is likely redundant to use both models. Consequently we used a single nodecov() model to analyse the empirical dataset.
2. Small sample sizes generate potential issues when relying on statistical significance from each individual network. Our simulations demonstrate both constraints on power to detect true effects and also elevated false positive rates that are not caused by biased parameter estimates. Individual movement networks such as these often have both small node sets and are sparse (low density of connections or low variance in edge weights). These results indicate the value of using meta-analysis type approaches to combine estimates from multiple individual movement networks in these situations as we did in our empirical study. Combining model outputs in this way mitigates issues related to false positive or false negative error rates when modelling the movement of any single individual while improving confidence of effects that are small or have large confidence intervals for any one individual but occur in a consistent direction between individuals.

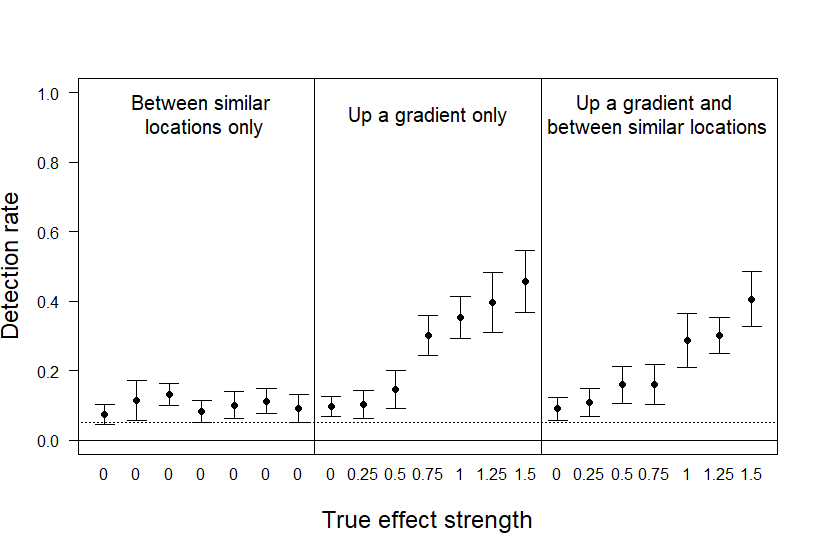

Fig. S2. Rate of detection of statistically significant results for the effect of factorA on the number of movements into a receiver location (nodeicov) in simulated movement networks with individuals initially located at random locations.

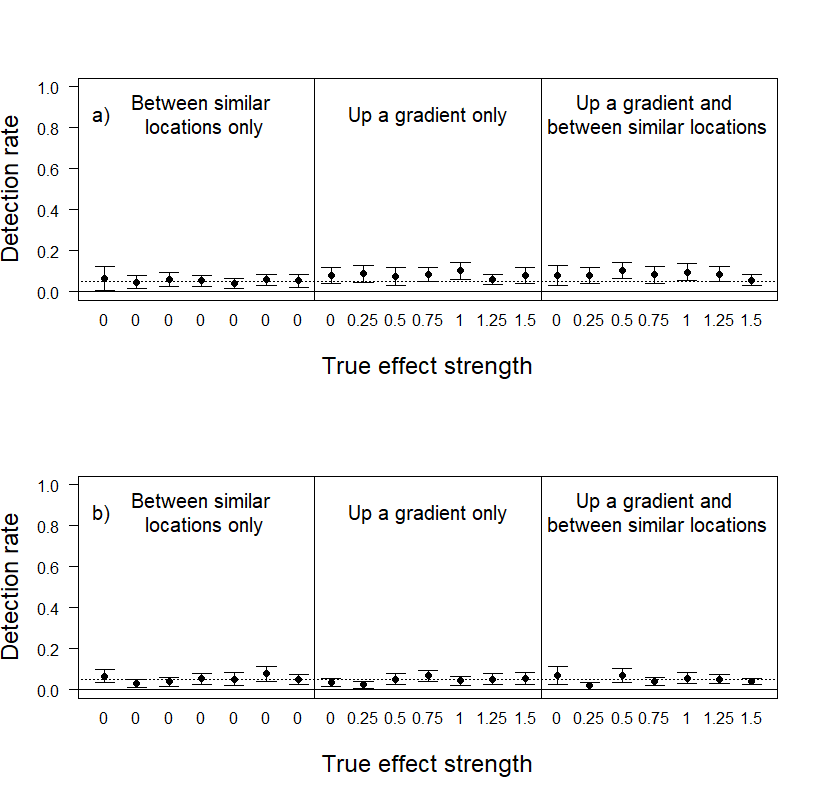

Fig. S3. Rate of detection of statistically significant results for the effect of a) factorB and b) factorC on the number of movements into a receiver location (nodeicov) in simulated movement networks with individuals initially located at random locations.

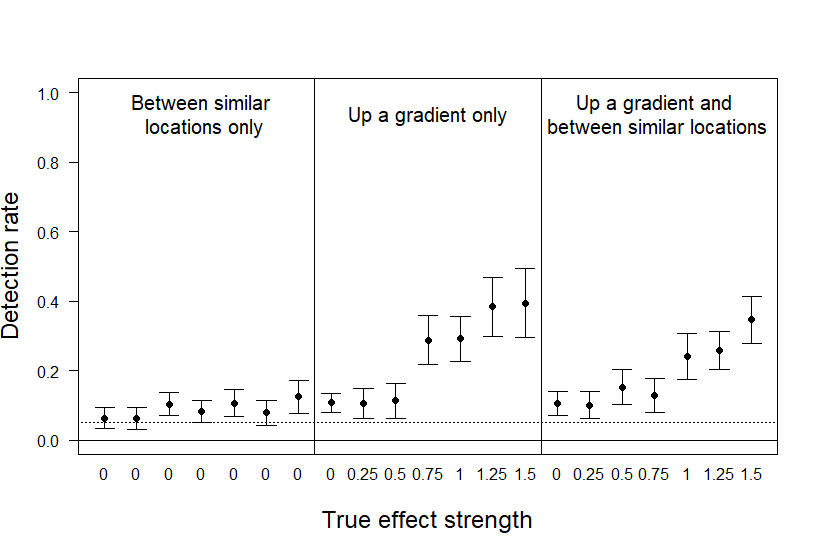

Fig. S4. Rate of detection of statistically significant results for the effect of factorA on the number of movements out from a receiver location (nodeocov) in simulated movement networks with individuals initially located at random locations.

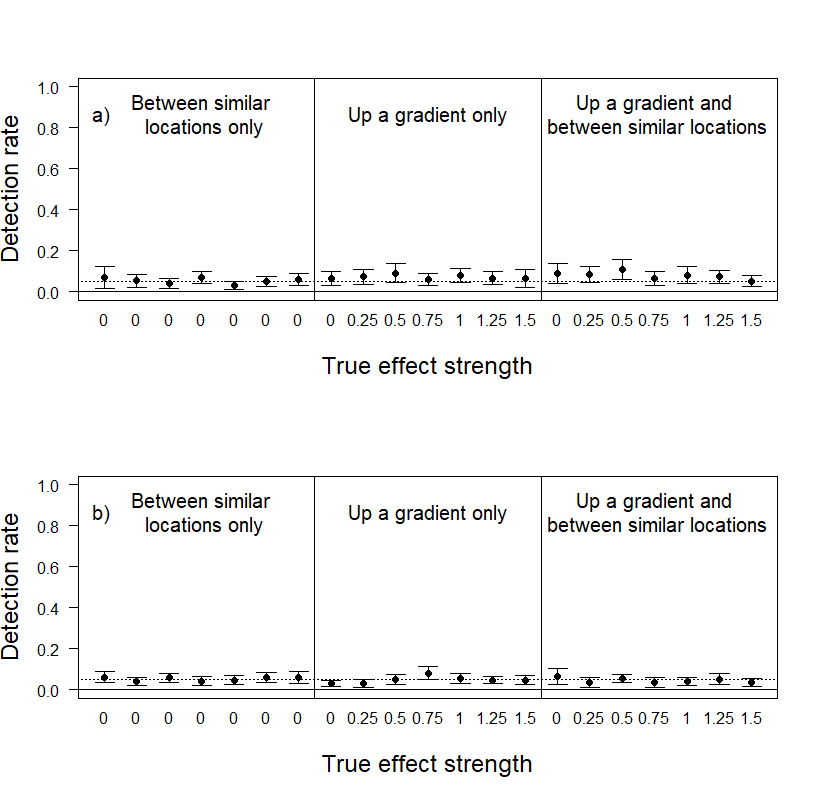

Fig. S5. Rate of detection of statistically significant results for the effect of a) factorB and b) factorC on the number of movements out from a receiver location (nodeocov) in simulated movement networks with individuals initially located at random locations.

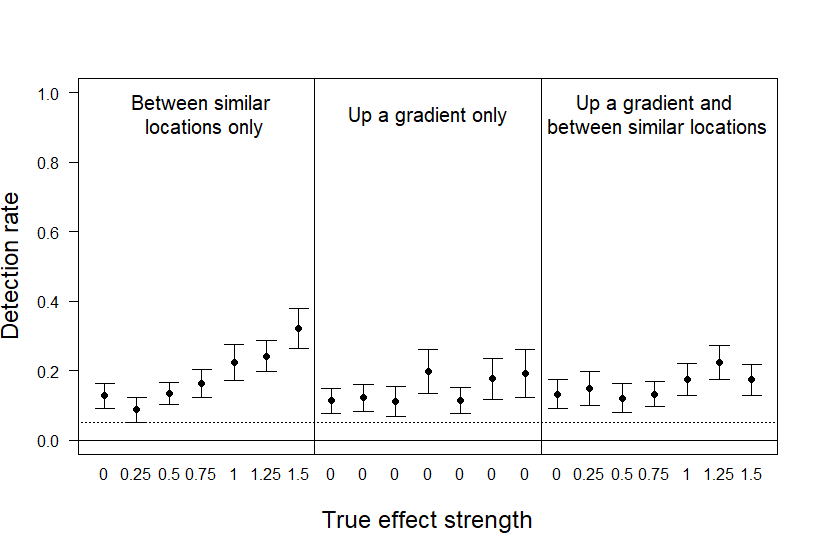

Fig. S6. Rate of detection of statistically significant results for the effect of similarity in factorA on the number of movements between receiver locations (absdiff) in simulated movement networks with individuals initially located at random locations.

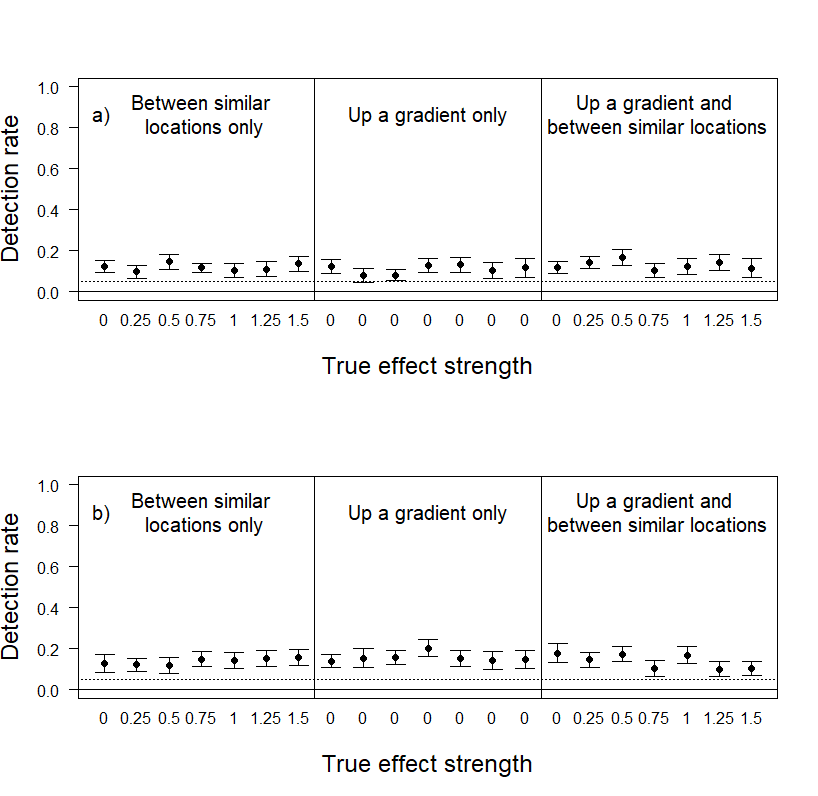

Fig. S7. Rate of detection of statistically significant results for the effect of similarity in a) factorB and b) factorC on the number of movements between receiver locations (absdiff) in simulated movement networks with individuals initially located at random locations.

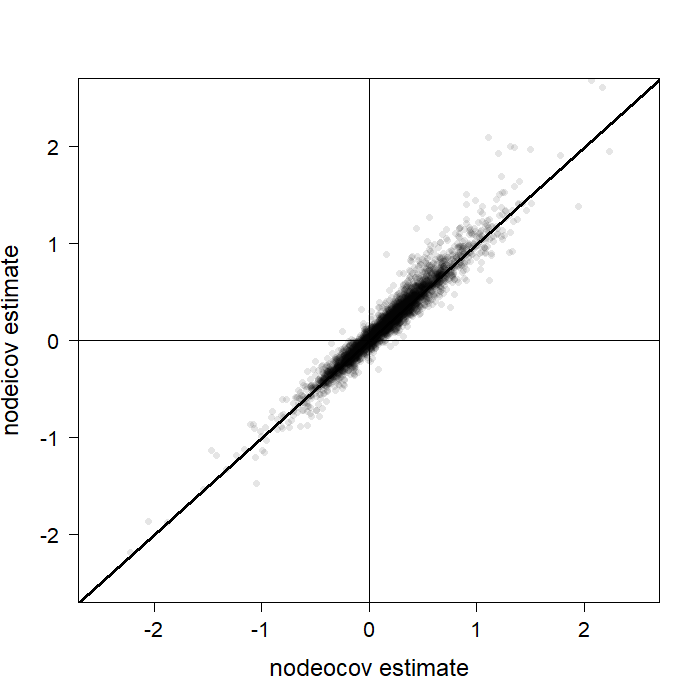

Fig. S8. The correlation between effect size estimates for factorA between nodeocov and nodeicov models in simulated movement networks (weighted networks only) with individuals initially located at random locations. 10 points lie outside the region plotted here and have been excluded for visualization purposes.

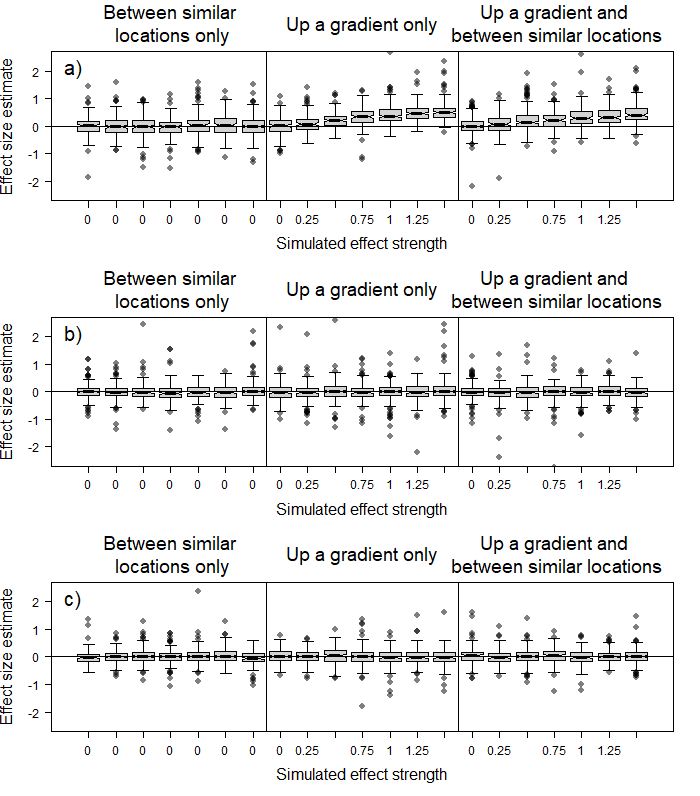

Fig. S9. Estimated effect size distributions for the effect of a) factorA, b) factorB and c) factorC on the number of movements into a receiver location (nodeicov) in simulated movement networks (weighted networks only) with individuals initially located at random locations. Solid lines represent the median and boxes the interquartile range. Whiskers extend to 1.5x the interquartile range. Some outlying points are not depicted to show the main results more clearly.

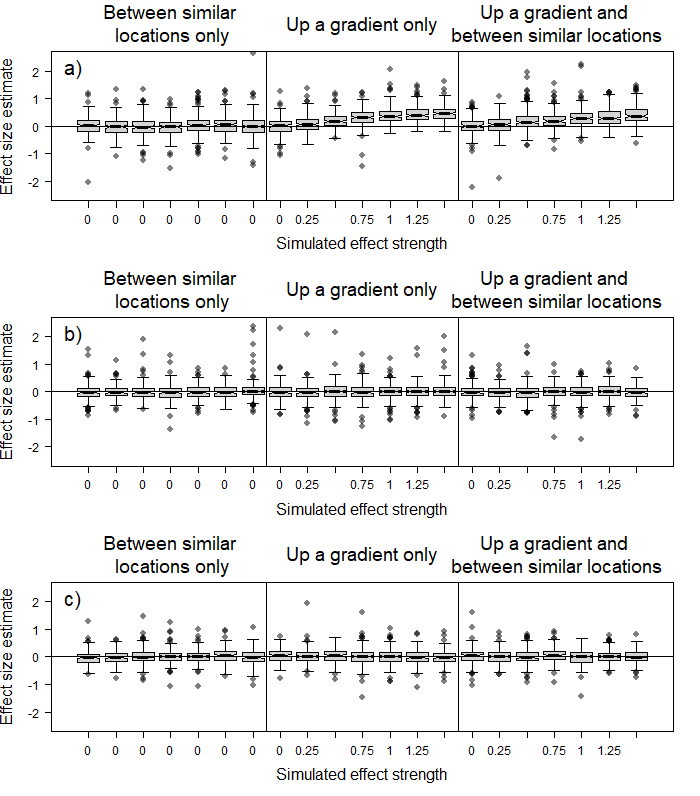

Fig. S10. Estimated effect size distributions for the effect of a) factorA, b) factorB and c) factorC on the number of movements out from a receiver location (nodeocov) in simulated movement networks (weighted networks only) with individuals initially located at random locations. Solid lines represent the median and boxes the interquartile range. Whiskers extend to 1.5x the interquartile range. Some outlying points are not depicted to show the main results more clearly.

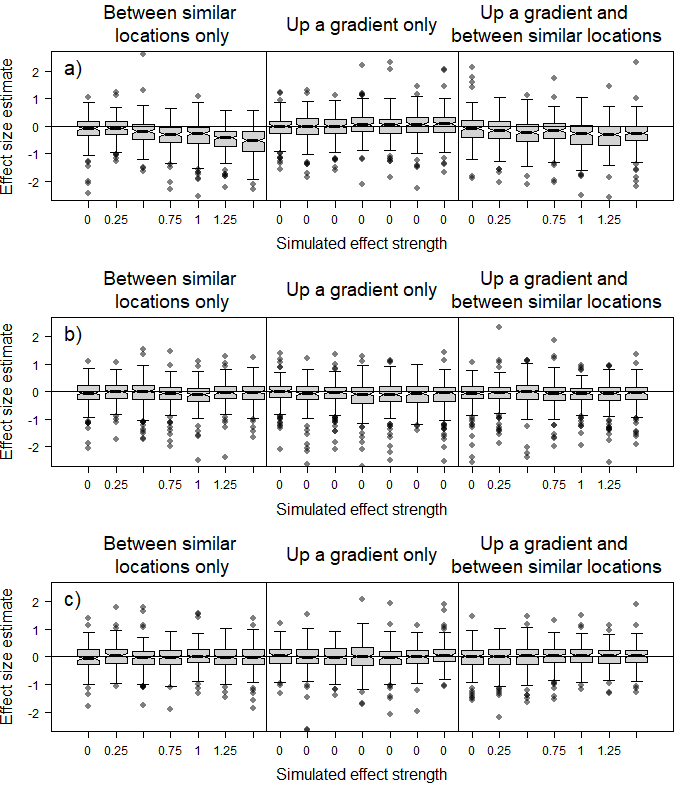

Fig. S11. Estimated effect size distributions for the effect of similarity in a) factorA, b) factorB and c) factorC on the number of movements between receiver locations (absdiff) in simulated movement networks (weighted networks only) with individuals initially located at random locations. Solid lines represent the median and boxes the interquartile range. Whiskers extend to 1.5x the interquartile range. Some outlying points are not depicted to show the main results more clearly.

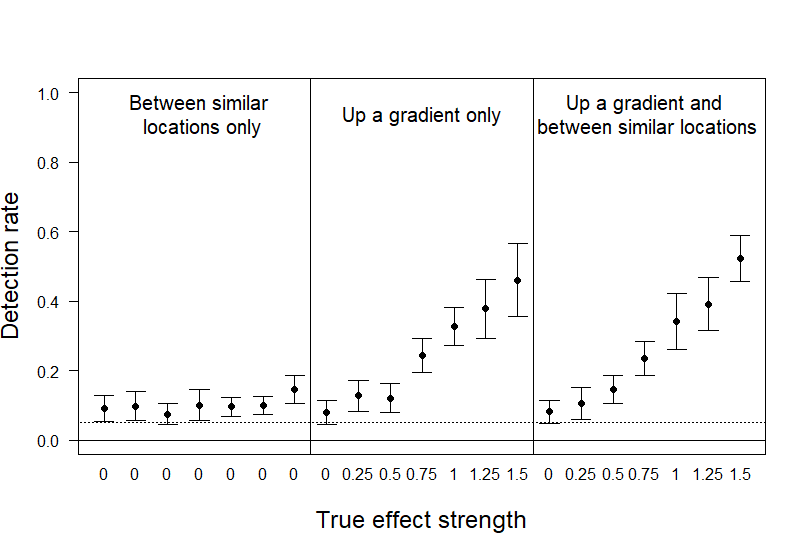

Fig. S12. Rate of detection of statistically significant results for the effect of factorA on the number of movements into a receiver location (nodeicov) in simulated movement networks with individual starting locations biased towards low factorA receivers.

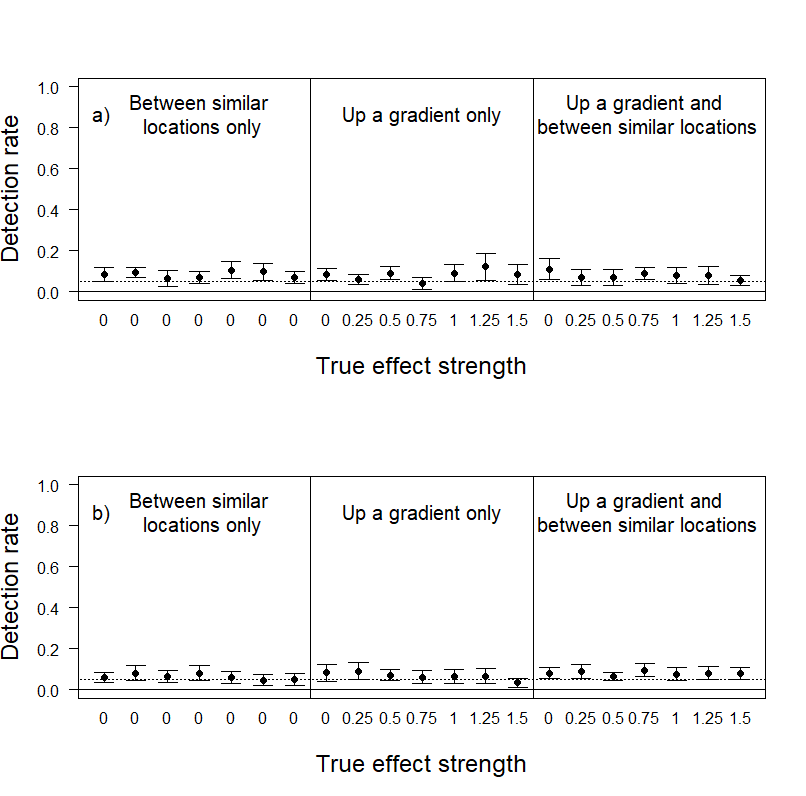

Fig. S13. Rate of detection of statistically significant results for the effect of a) factorB and b) factorC on the number of movements into a receiver location (nodeicov) in simulated movement networks with individual starting locations biased towards low factorA receivers.

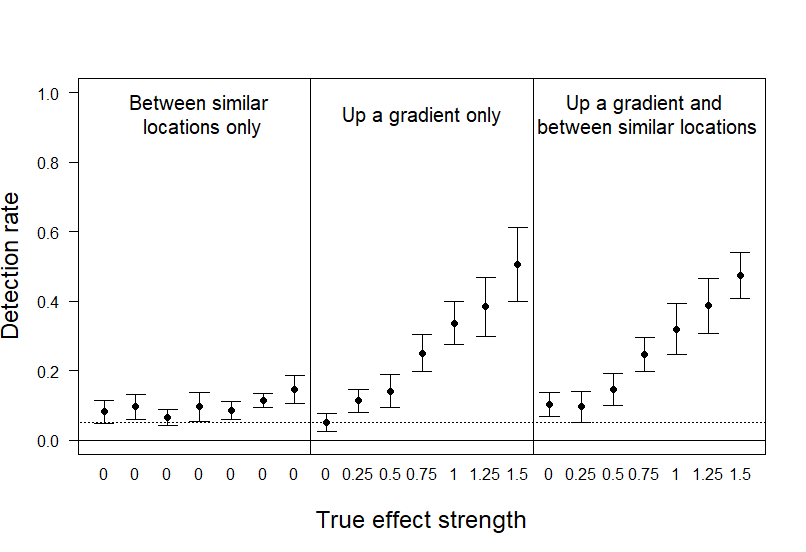

Fig. S14. Rate of detection of statistically significant results for the effect of factorA on the number of movements out from a receiver location (nodeocov) in simulated movement networks with individual starting locations biased towards low factorA receivers.

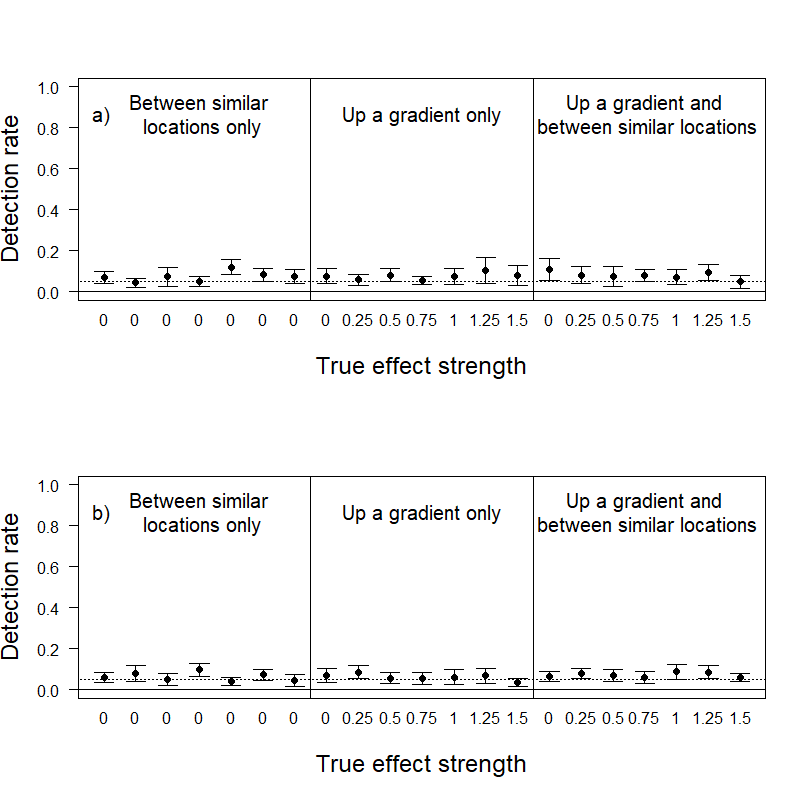

Fig. S15. Rate of detection of statistically significant results for the effect of a) factorB and b) factorC on the number of movements out from a receiver location (nodeocov) in simulated movement networks with individual starting locations biased towards low factorA receivers.

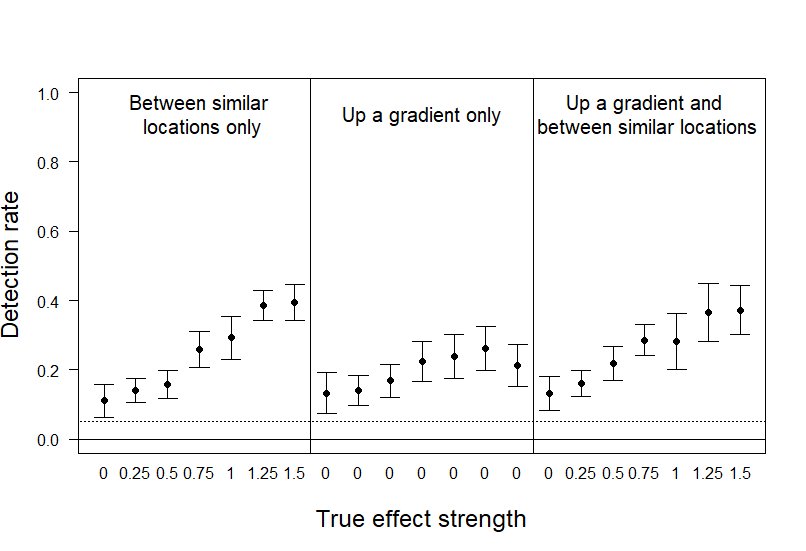

Fig. S16. Rate of detection of statistically significant results for the effect of similarity in factorA on the number of movements between receiver locations (absdiff) in simulated movement networks with individual starting locations biased towards low factorA receivers.

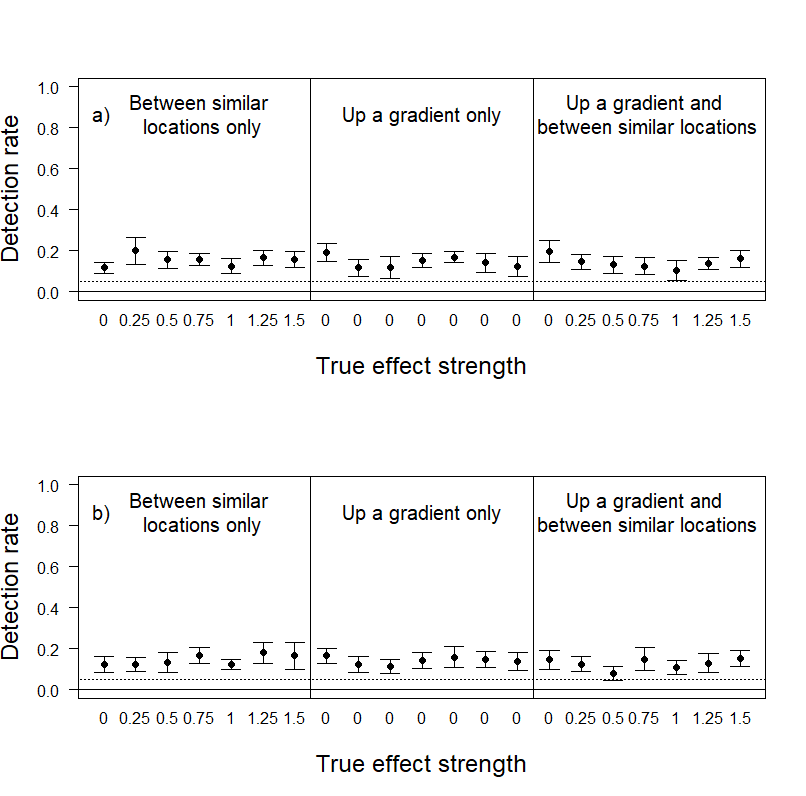

Fig. S17. Rate of detection of statistically significant results for the effect of similarity in a) factorB and b) factorC on the number of movements between receiver locations (absdiff) in simulated movement networks with individual starting locations biased towards low factorA receivers.

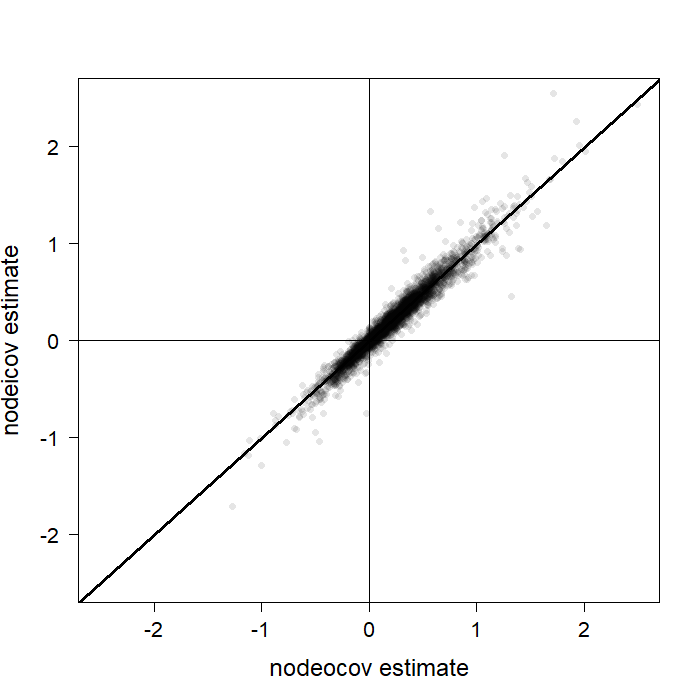

Fig. S18. The correlation between effect size estimates for factorA between nodeocov and nodeicov models in simulated movement networks (weighted networks only) with individual starting locations biased towards low factorA receivers. 5 points lie outside the region plotted here and have been excluded for visualization purposes.

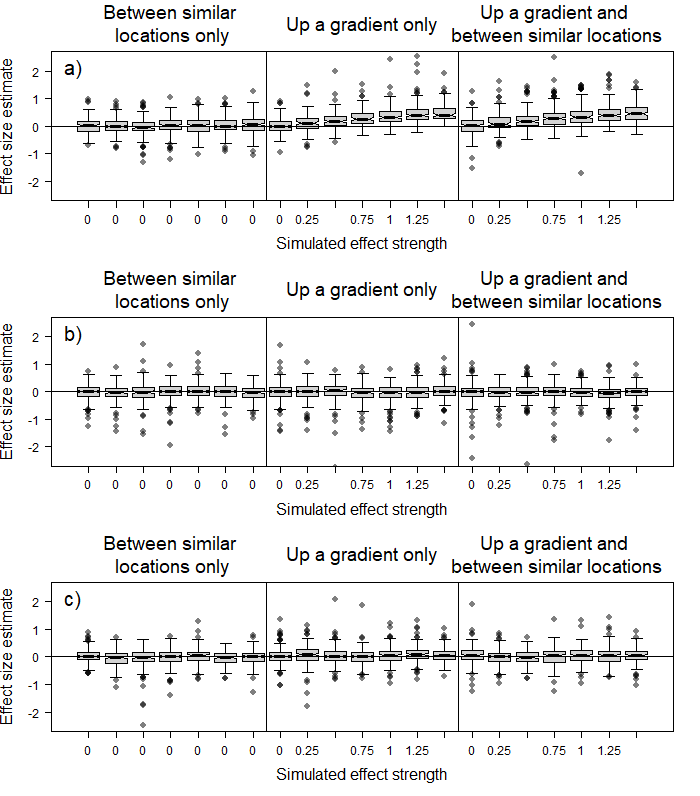

Fig. S19. Estimated effect size distributions for the effect of a) factorA, b) factorB and c) factorC on the number of movements into a receiver location (nodeicov) in simulated movement networks (weighted networks only) with individual starting locations biased towards low factorA receivers. Solid lines represent the median and boxes the interquartile range. Whiskers extend to 1.5x the interquartile range. Some outlying points are not depicted to show the main results more clearly.

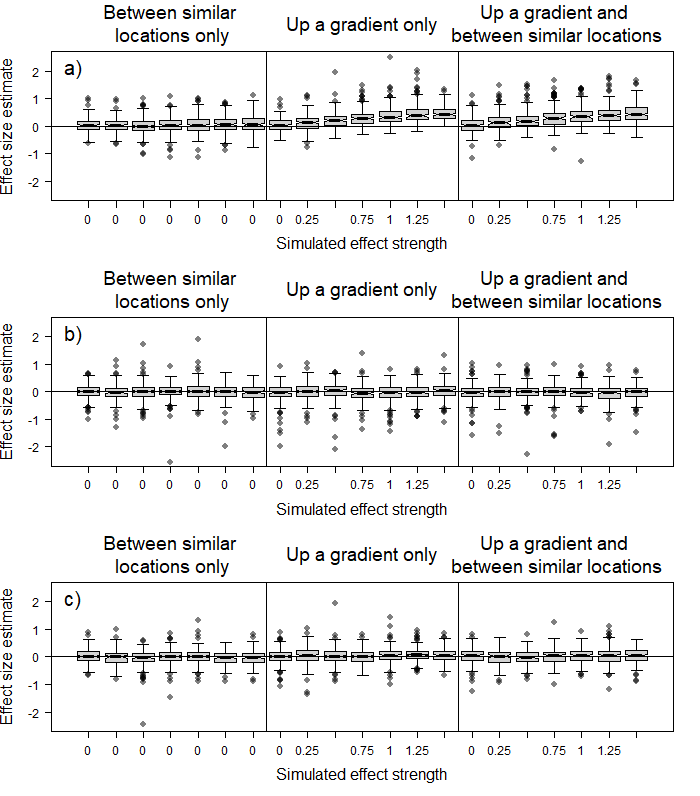

Fig. S20. Estimated effect size distributions for the effect of a) factorA, b) factorB and c) factorC on the number of movements out from a receiver location (nodeocov) in simulated movement networks (weighted networks only) with individual starting locations biased towards low factorA receivers. Solid lines represent the median and boxes the interquartile range. Whiskers extend to 1.5x the interquartile range. Some outlying points are not depicted to show the main results more clearly.

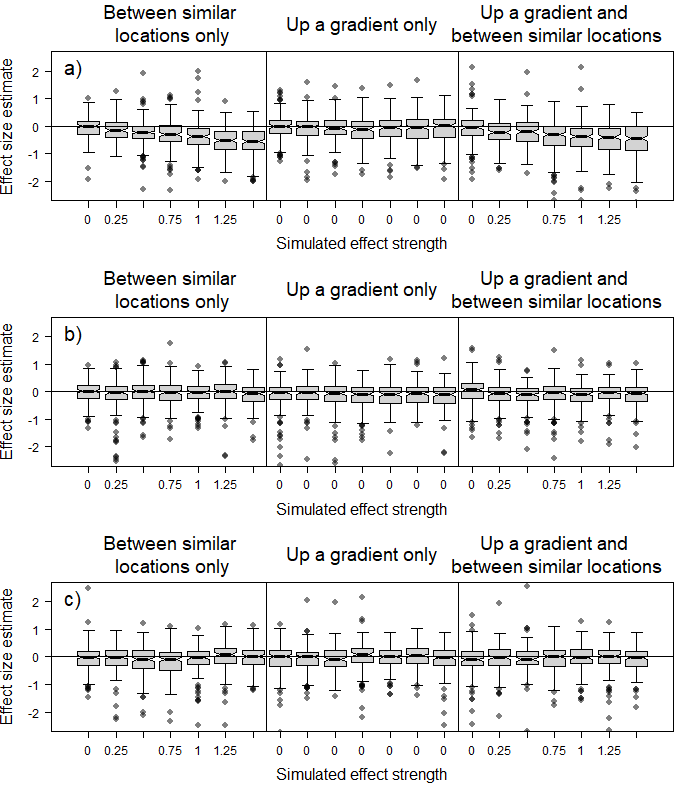

Fig. S21. Estimated effect size distributions for the effect of similarity in a) factorA, b) factorB and c) factorC on the number of movements between receiver locations (absdiff) in simulated movement networks (weighted networks only) with individual starting locations biased towards low factorA receivers. Solid lines represent the median and boxes the interquartile range. Whiskers extend to 1.5x the interquartile range. Some outlying points are not depicted to show the main results more clearly.

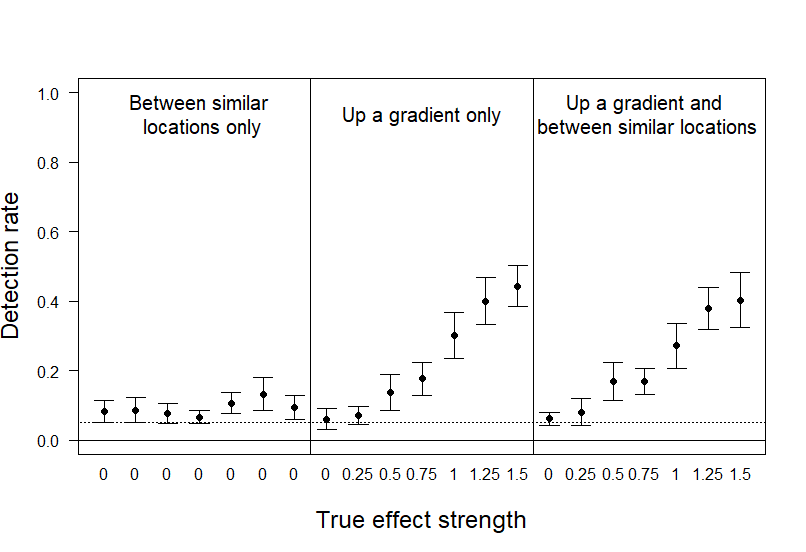

Fig. S22. Rate of detection of statistically significant results for the effect of factorA on the number of movements into a receiver location (nodeicov) in simulated movement networks with individual starting locations biased towards high factorA receivers.

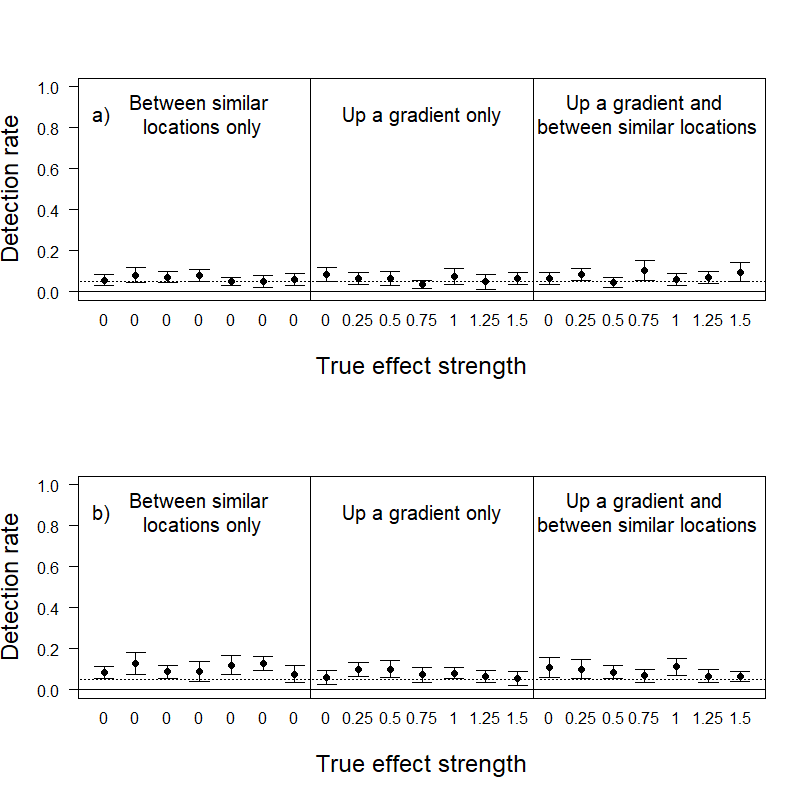

Fig. S23. Rate of detection of statistically significant results for the effect of a) factorB and b) factorC on the number of movements into a receiver location (nodeicov) in simulated movement networks with individual starting locations biased towards high factorA receivers.

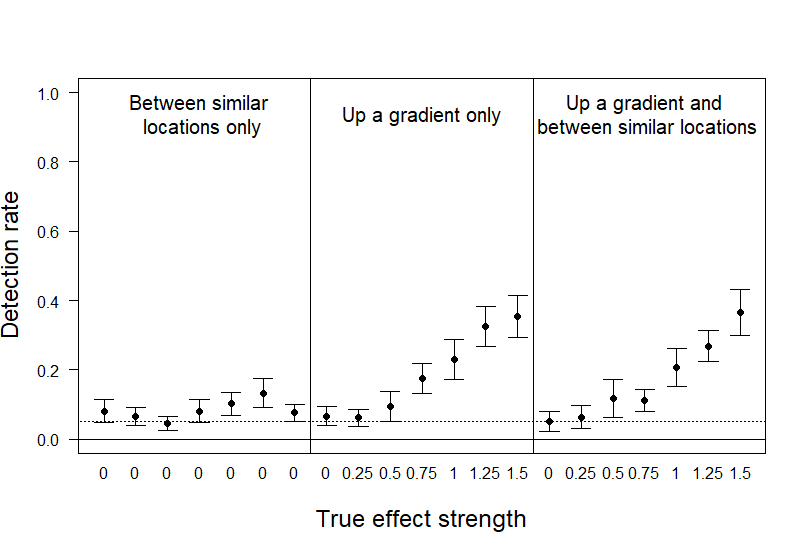

Fig. S24. Rate of detection of statistically significant results for the effect of factorA on the number of movements out from a receiver location (nodeocov) in simulated movement networks with individual starting locations biased towards high factorA receivers.

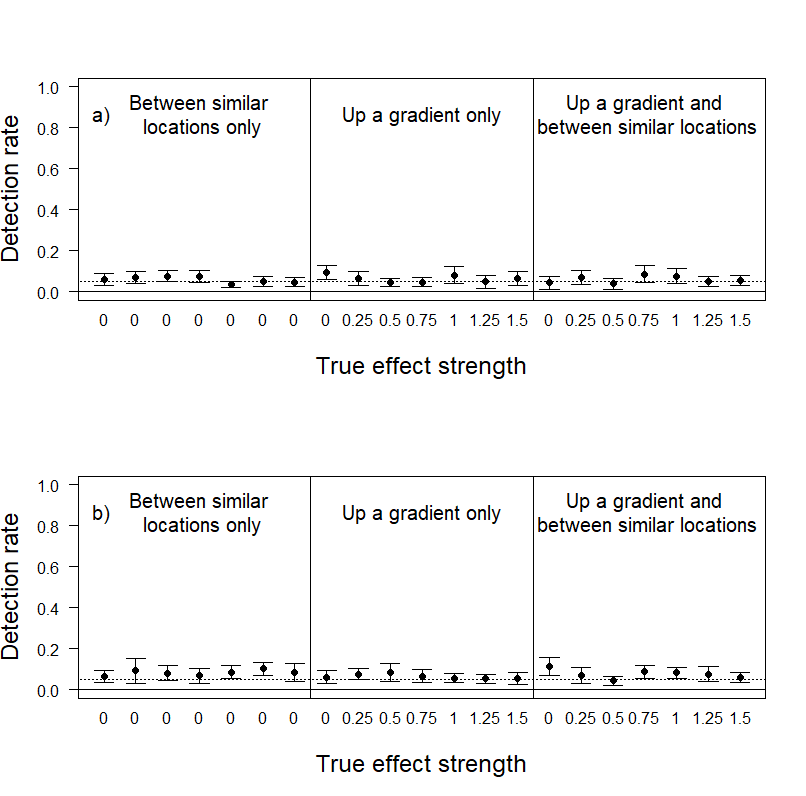

Fig. S25. Rate of detection of statistically significant results for the effect of a) factorB and b) factorC on the number of movements out from a receiver location (nodeocov) in simulated movement networks with individual starting locations biased towards high factorA receivers.

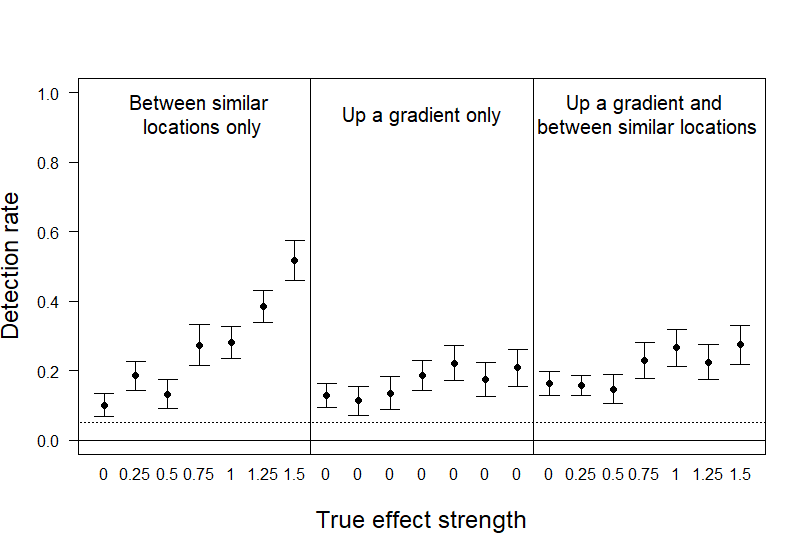

Fig. S26. Rate of detection of statistically significant results for the effect of similarity in factorA on the number of movements between receiver locations (absdiff) in simulated movement networks with individual starting locations biased towards high factorA receivers.

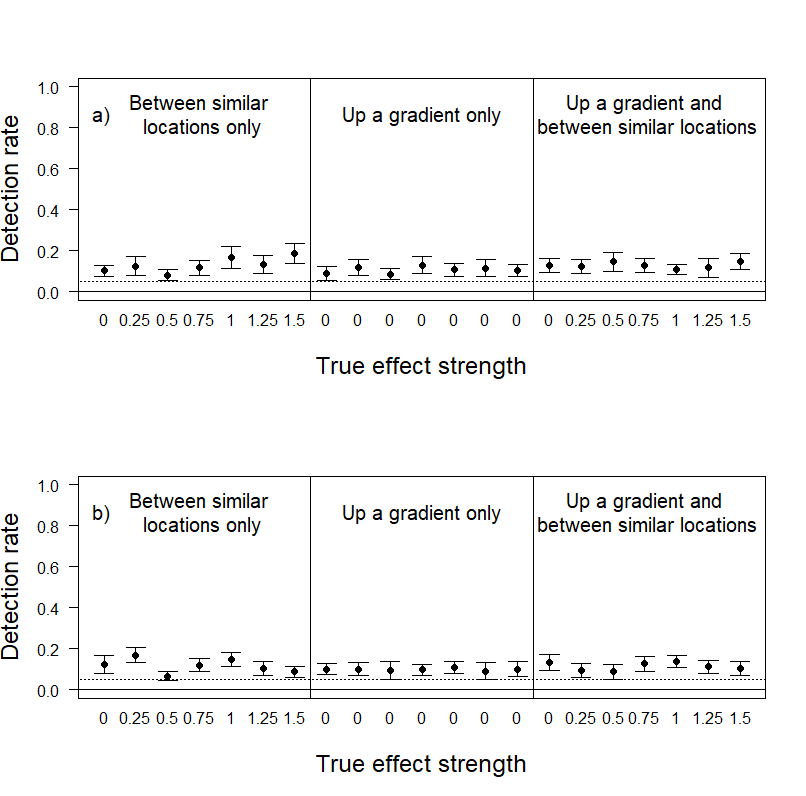

Fig. S27. Rate of detection of statistically significant results for the effect of similarity in a) factorB and b) factorC on the number of movements between receiver locations (absdiff) in simulated movement networks with individual starting locations biased towards high factorA receivers.

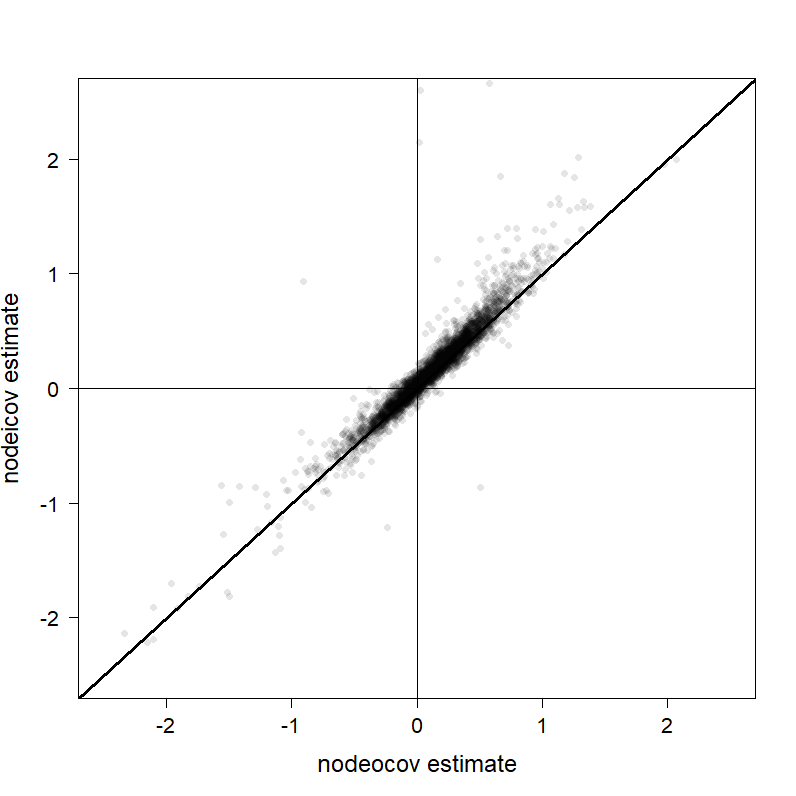

Fig. S28. The correlation between effect size estimates for factorA between nodeocov and nodeicov models in simulated movement networks (weighted networks only) with individual starting locations biased towards high factorA receivers. 14 points lie outside the region plotted here and have been excluded for visualization purposes.

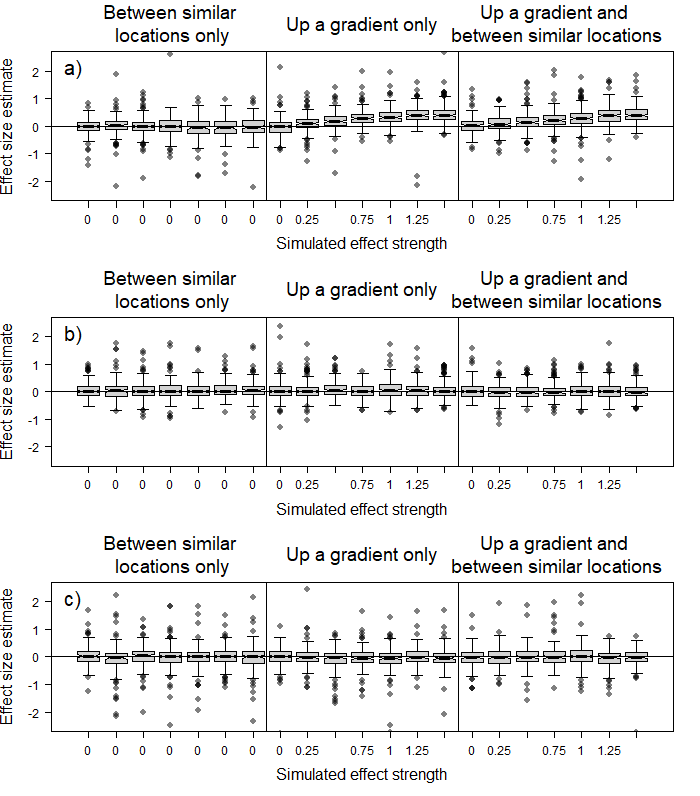

Fig. S29. Estimated effect size distributions for the effect of a) factorA, b) factorB and c) factorC on the number of movements into a receiver location (nodeicov) in simulated movement networks (weighted networks only) with individual starting locations biased towards high factorA receivers. Solid lines represent the median and boxes the interquartile range. Whiskers extend to 1.5x the interquartile range. Some outlying points are not depicted to show the main results more clearly.

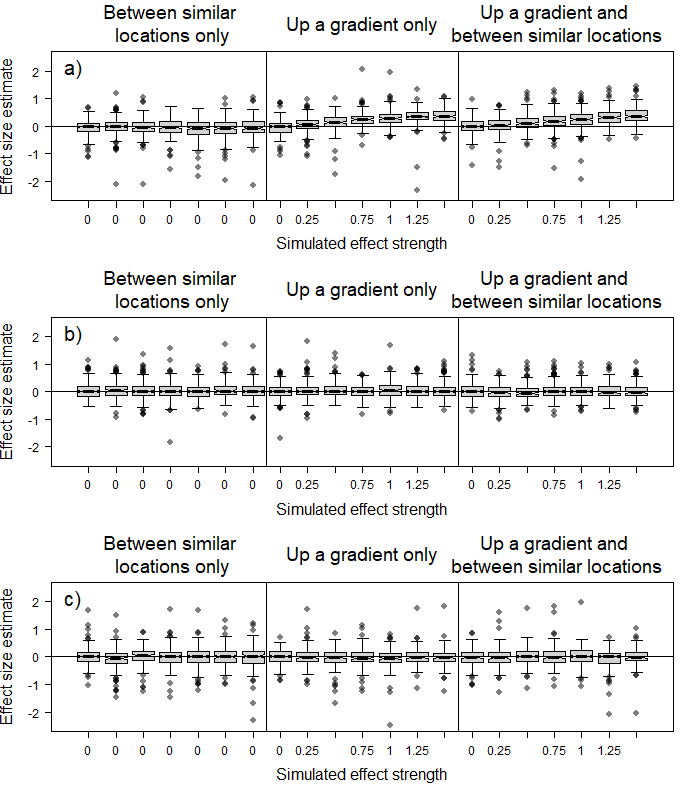

Fig. S30. Estimated effect size distributions for the effect of a) factorA, b) factorB and c) factorC on the number of movements out from a receiver location (nodeocov) in simulated movement networks (weighted network only) with individual starting locations biased towards high factorA receivers. Solid lines represent the median and boxes the interquartile range. Whiskers extend to 1.5x the interquartile range. Some outlying points are not depicted to show the main results more clearly.

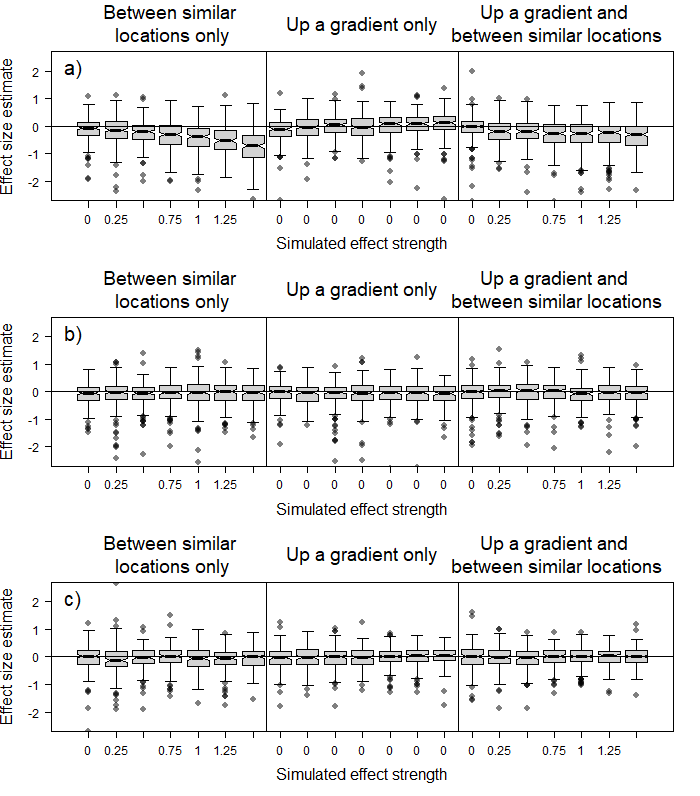

Fig. S31. Estimated effect size distributions for the effect of similarity in a) factorA, b) factorB and c) factorC on the number of movements between receiver locations (absdiff) in simulated movement networks (weighted networks only) with individual starting locations biased towards high factorA receivers. Solid lines represent the median and boxes the interquartile range. Whiskers extend to 1.5x the interquartile range. Some outlying points are not depicted to show the main results more clearly.
